# Dimensionality of genomic information and its impact on GWA and variant selection: a simulation study

**DOI:** 10.1101/2022.04.13.488175

**Authors:** Sungbong Jang, Shogo Tsuruta, Natalia Galoro Leite, Ignacy Misztal, Daniela Lourenco

## Abstract

**Background:** Identifying true-positive variants in genome-wide associations (GWA) depends on several factors, including the number of genotyped individuals. The limited dimensionality of the genomic information may give insights into the optimal number of individuals to use in GWA. This study investigated different discovery set sizes in GWA based on the number of largest eigenvalues explaining a certain proportion of variance in the genomic relationship matrix (**G**). An additional investigation included the change in accuracy by adding variants, selected based on different set sizes, to the regular SNP chips used for genomic prediction.

**Methods:** Sequence data were simulated containing 500k SNP with 200 or 2000 quantitative trait nucleotides (QTN). A regular 50k panel included one every ten simulated SNP. Effective population size (*Ne*) was 20 and 200. The GWA was performed with the number of genotyped animals equivalent to the number of largest eigenvalues of **G** (EIG) explaining 50, 60, 70, 80, 90, 95, 98, and 99% of the variance. In addition, the largest discovery set consisted of 30k genotyped animals. Limited or extensive phenotypic information was mimicked by changing the trait heritability. Significant and high effect size SNP were added to the 50k panel and used for single-step GBLUP with and without weights.

**Results:** Using the number of genotyped animals corresponding to at least EIG98 enabled the identification of QTN with the largest effect sizes when *Ne* was large. Smaller populations required more than EIG98. Furthermore, using genotyped animals with higher reliability (i.e., higher trait heritability) helped better identify the most informative QTN. The greatest prediction accuracy was obtained when the significant or the high effect SNP representing twice the number of simulated QTN were added to the 50k panel. Weighting SNP differently did not increase prediction accuracy, mainly because of the size of the genotyped population.

**Conclusions:** Accurately identifying causative variants from sequence data depends on the effective population size and, therefore, the dimensionality of genomic information. This dimensionality can help identify the suitable sample size for GWA and could be considered for variant selection. Even when variants are accurately identified, their inclusion in prediction models has limited implications.

## Background

Several factors influence the statistical power to identify causative variants in genome-wide associations (GWA), including the number of quantitative trait nucleotides (QTN) affecting the trait, the number of single nucleotide polymorphisms (SNP) in the discovery panel, the number of genotyped individuals [1], and the size of the genome blocks segregating in the population [2], among others. Those genome blocks are chromosome segments inherited from founders and are subject to recombination every generation. Stam [3] showed that segments are of different sizes but with a mean size of 1/4*N*_*e*_, where *N*_*e*_ is the effective population size. Given a species with a genome length equal to *L* Morgans, the number of independent chromosome segments (*M*_*e*_) segregating in a population can be calculated as 4*N*_*e*_*L*.

Animal populations have lower *N*_*e*_ than human populations, implying smaller *M*_*e*_. Pocrnic et al. [4] showed that although millions of individuals can be genotyped, non-redundant information is finite, which means the genomic information has a limited dimensionality; therefore, the additive genetic information in a population is contained in a limited *M*_*e*_. The same authors related the limited dimensionality to *M*_*e*_ = 4*N*_*e*_*L* and observed that this quantity corresponds to the number of largest eigenvalues explaining 98% (EIG98) of the variance of the genomic relationship matrix (**G**). In cattle populations, EIG98 varies from 10K to 14K and is about 4K in pigs and chickens. The minimum number of SNP needed to cover those segments is approximately 12*M*_*e*_ [5].

With the availability of sequence information, causal variants are expected to be in the data, generating more opportunities for discovery than with mid-density SNP panels [6]. When the causal variants are known and included in the usual SNP panels, accuracy of predicting genomic estimated breeding values (GEBV) should increase. This is clearly observed in simulated studies where the QTN and their effects are known [7, 8]. However, the accuracy increased by using significant variants from the sequence in real populations is almost inexistent [9-11]. This raises a question on the effectiveness of GWA in real populations. Although most traits of economic importance in farm animal populations are polygenic, very few peaks are usually statistically associated with traits of interest.

Misztal et al. [12] investigated the distribution of SNP around the QTN and the ability to identify QTN depending on the *N*_*e*_ in simulated populations. They found that identifying QTN in populations with small *N*_*e*_ (i.e., 60) required three times more genotyped animals with phenotypes than in populations with large *N*_*e*_ (i.e., 600). However, not all simulated QTN were identified, independently of the *N*_*e*_ or amount of data. Distinguishing between noise and the true signal is more difficult in small populations because of longer chromosome segments and the uncertainty about the exact QTN location. Additionally, the level of noise may mask the signal, preventing associations. With sequence data, a clear GWA resolution for small populations may be even harder to achieve due to the reasons mentioned above.

Although it is well-known that increasing the sample size for GWA improves the resolution, the links among the number of genotyped individuals, *N*_*e*_, *M*_*e*_, and GWA resolution are missing. Additionally, understanding the appropriate sample size for variant discovery, especially with sequence data, can help to alleviate both the economic and computational costs for practical applications. Based on the limited dimensionality of the genomic information, there may be an optimal number of animals that carry all the independent chromosome segments segregating in the population, and consequently, all the genomic information available in the population [4]. When animals have lots of information, GEBV are estimated with high accuracy. Knowing that GEBV can be backsolved to SNP effects raises the question on whether GWA resolution is high when *M*_*e*_ animals with high accuracy GEBV are used. Therefore, we hypothesize that the ability to identify causative variants is high when the sample size for GWA approaches *M*_*e*,_ and using a larger sample size may not further improve GWA resolution. Here we used the number of eigenvalues explaining different proportions of the variance in **G** to assess the dimensionality of the genomic information and used this number as the sample size in GWA. We used simulated populations with varying *N*_*e*_, number of QTN, and amount of information on genotyped individuals. We also evaluated the impact of incorporating the pre-selected variants, from GWA with different sample sizes based on dimensionality, to a 50k SNP chip for genomic prediction using weighted and unweighted single-step GBLUP.

## Methods

### Data simulation

QMSim [13] was used to simulate a quantitative trait with 0.3, 0.9, and 0.99 heritability. Different heritabilities mimicked limited or extensive phenotypic information. The historical population was simulated for 2,000 non-overlapping generations with an increase in size from 1,000 (generation -2,000) to 50,000 (generation -1,000), and a decrease from 50,000 (generation -999) to 20,000 (generation 0) to create LD and mutation−drift equilibrium. Random mating and no selection or migration were assumed in the historical population. Recent populations of *N*_*e*_ equal 20 (*N*_*e*_20) and 200 (*N*_*e*_200) were simulated by changing the number of breeding males from 5 to 50 but keeping the number of females at 15,000. The founders of the recent populations came from generation 0 of the historical population. Twenty generations of random mating were carried out, considering a replacement rate of 80% for sires and 30% for dams. Animals were randomly selected and culled based on age. A total of 315,005 and 315,050 animals were generated in the recent population for *N*_*e*_20 and *N*_*e*_200, respectively. However, only animals from generations 11−20 had phenotypic and pedigree information that was used for the current study. Of those, 75,000 animals from generations 16-20 were genotyped (N = 15,000 in each generation). The phenotype was the sum of an overall mean equal to 1.0, true breeding value (TBV), and random residual effect. The phenotypic variance was set to 1.0, whereas the additive genetic variance was 0.3, 0.9, or 0.99, all explained by the simulated QTN.

To mimic the bovine genome, we simulated 29 chromosomes with a total length of 23.19 Morgans. The overall number of SNP was 500,000, all with minor allele frequency greater than 0.05, whereas QTN numbers were 200 and 2,000 for the scenarios Q200 and Q2000, respectively. Biallelic SNP and QTN were randomly placed on each chromosome, with numbers varying from 9,000−35,000 SNP and 8−31 (Q200) or 80−320 (Q2000) QTN. The QTN effects were sampled from a gamma distribution with shape parameter 0.4 and scale parameter calculated internally for a genetic variance of 0.3, 0.9, and 0.99, depending on the scenario. A recurrent mutation rate of 2.5 × 10^−5^ was assumed for both SNP and QTN. A regular 50k panel was created for genomic predictions (GP) that included one every ten simulated SNP. Because different simulation replicates would generate different QTN positions and effects, no replicate was used to obtain consistent GWA results.

### Genotype scenarios – heritability and the presence of QTN in the data

Trait heritabilities of 0.3, 0.9, and 0.99 were simulated to represent the animals with low reliability of EBV (H30), high reliability of EBV (H90), and very high reliability of EBV (H99), respectively. Therefore, higher heritabilities mean more information was added to the simulated animals without directly changing the number of records assigned to them [14]. Further, sequence data scenarios were created after the simulation. We assumed that QTN were in the data (withQTN), following the general assumption for sequence data; therefore, the SNP and QTN files were combined based on the corresponding maps. Descriptions for all the scenarios and combinations are in Table 1.

**Table 1.** Description of all GWA scenarios.

| Scenario description | Ne | Number of QTN | Heritability |
| --- | --- | --- | --- |
| Ne20 Q200 H30 | 20 | 200 | 0.3 |
| Ne20 Q200 H90 | 20 | 200 | 0.9 |
| Ne20 Q200 H99 | 20 | 200 | 0.99 |
| Ne20 Q2000 H30 | 20 | 2000 | 0.3 |
| Ne20 Q2000 H90 | 20 | 2000 | 0.9 |
| Ne20 Q2000 H99 | 20 | 2000 | 0.99 |
| Ne200 Q200 H30 | 200 | 200 | 0.3 |
| Ne200 Q200 H90 | 200 | 200 | 0.9 |
| Ne200 Q200 H99 | 200 | 200 | 0.99 |
| Ne200 Q2000 H30 | 200 | 2000 | 0.3 |
| Ne200 Q2000 H90 | 200 | 2000 | 0.9 |
| Ne200 Q2000 H99 | 200 | 2000 | 0.99 |

### Discovery, training, and test sets

Before the GWA analyses, all genotyped animals were separated into three non-overlapping data sets: discovery, training, and test. The test set was composed of genotyped animals from the last generation (N = 15,000), whereas the remaining genotyped animals (N = 60,000) were randomly assigned to the discovery and training sets (N = 30,000, respectively). To test the possible bias in GP by using the same data set for discovery and training, two different schemes were designed: 1) discovery = training: genotyped animals used for discovery were further used for training, 2) discovery ≠ training: a different set of genotyped animals were used for discovery and training.

### EIG*x* scenarios for discovery and training

Different scenarios were made based on the dimensionality of the genomic information to investigate the effect of sample size for discovery and training. The number of genotyped animals in each discovery and training set (EIG*x*) was equivalent to the number of largest eigenvalues explaining *x* percent of the variance in **G**, where *x* assumed the values 50, 60, 70, 80, 90, 95, 98, or 99. For example, the number of largest eigenvalues explaining 50% of the variance in **G** was 530 in the *N*_*e*_200 Q2000 H30 scenario (Table 2); thus, the size of discovery and training sets in scenario EIG50 was set to 530. In addition, one extra scenario (ALL) in which the discovery and training sets consisted of all available genotyped animals (N = 30,000) was also evaluated. The number of largest eigenvalues explaining *x* percent (50, 60, 70, 80, 90, 95, 98, 99) of the variance in **G** was computed by squaring the singular values from the matrix of genotypes centered for current allele frequencies (**M**), using all the simulated genotyped animals (N = 75,000). The singular value decomposition was done in preGSf90 [15]. For that, **G** was constructed using all simulated SNP without QTN. All genotyped animals for each discovery and training set were randomly selected beginning from the scenario explaining the least proportion of variance (EIG50). Ensuring consistent results involved keeping all the animals from a previous scenario when moving to the next one, e.g., genotyped animals in EIG60 contained the ones from EIG50. The number of genotyped animals for all scenarios used as discovery and training sets is described in Table 2.

**Table 2.** Number of genotyped animals for all scenarios in both discovery and training sets.

|  | Ne20 | Ne20 | Ne20 | Ne20 | Ne20 | Ne20 | Ne200 | Ne200 | Ne200 | Ne200 | Ne200 | Ne200 |
| --- | --- | --- | --- | --- | --- | --- | --- | --- | --- | --- | --- | --- |
|  | Q200 H30 | Q200 H90 | Q200 H99 | Q2000 H30 | Q2000 H90 | Q2000 H99 | Q200 H30 | Q200 H90 | Q200 H99 | Q2000 H30 | Q2000 H90 | Q2000 H99 |
| EIG50 | 80 | 80 | 80 | 80 | 80 | 80 | 510 | 520 | 550 | 530 | 500 | 510 |
| EIG60 | 130 | 140 | 140 | 140 | 140 | 130 | 900 | 910 | 940 | 920 | 900 | 900 |
| EIG70 | 220 | 240 | 230 | 220 | 230 | 220 | 1,500 | 1,550 | 1,600 | 1,540 | 1,500 | 1500 |
| EIG80 | 390 | 410 | 410 | 400 | 400 | 400 | 2,600 | 2,650 | 2,700 | 2,650 | 2,600 | 2,600 |
| EIG90 | 860 | 900 | 890 | 890 | 880 | 840 | 5,100 | 5,250 | 5,300 | 5,300 | 5,100 | 5,100 |
| EIG95 | 1,700 | 1,770 | 1,800 | 1,800 | 1,750 | 1,700 | 8,600 | 8,800 | 8,800 | 8,800 | 8,600 | 8,600 |
| EIG98 | 4,000 | 4,100 | 4,100 | 4,100 | 4,100 | 4,000 | 15,000 | 15,200 | 15,200 | 15,200 | 15,000 | 15,000 |
| EIG99 | 6,900 | 7,100 | 7,100 | 7,100 | 7,000 | 6,900 | 21,000 | 22,000 | 21,400 | 22,000 | 22,000 | 21,000 |
| All | 30,000 | 30,000 | 30,000 | 30,000 | 30,000 | 30,000 | 30,000 | 30,000 | 30,000 | 30,000 | 30,000 | 30,000 |

### Preselection of variants (TOP*v* scenarios)

Different numbers of variants were selected from GWA to be included in the 50k SNP panel for GP. Each QTN scenario had a specific number of selected SNP based on the order of the p-values (TOP*v*) or statistical significance using Bonferroni corrected p-values (SIG). For Q200, *v* corresponded to 10, 50, 100, 200, and 400, whereas for Q2000 *v* assumed the values of 10, 100, 500, 1000, 2000, and 4000.

### Association among number of significant QTN, sample size, and EBV reliability

In the current study, we approximated sample size based on the total proportion of variance explained by significantly identified QTN from the results of different heritability scenarios. This approximation was done by local polynomial regression [16] using the ‘loess’ and ‘approx’ function in the R, and the resulting sample size was represented by SS_pol_.

The main purpose of comparing scenarios with different heritabilities was to investigate the effect of using genotyped animals with low to high reliability of EBV on GWA performance; however, not all the genotyped animals have high reliability of EBV. Therefore, in this study, we investigated the corresponding sample size from low to high heritability scenarios at the point where the same percentage of variance was explained; we estimated the sample size using H30 as a benchmark. This helped us identify how many samples are needed for GWA given the average reliability of breeding values for the animals in the population and the benchmark reliability. For that, derived an equation to estimate the sample size relating the total number of samples, *M*_*e*_, *N*_*e*_, proportion of additive genetic variance explained by significant QTN (*%Var*), and reliability of EBV as:

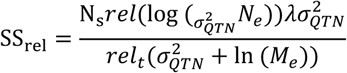

in which SS_rel_ is the approximated sample size for target reliability, N_s_ is the benchmark sample size, *rel* and *rel*_*t*_ are the benchmark and target reliabilities of EBV (heritability), *λ* is a constant equal to 0.4, and 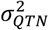 is the %var explained by identified QTN. The total proportion of genetic variance explained by the identified QTN was calculated as the sum of the genetic variance explained by each QTN. As QTN effects were given by the simulation, the percentage of genetic variance explained by individual QTN was calculated as:

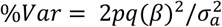

where the *p* and *q* are the major and minor allele frequency of the QTN, *β* is the QTN effect, 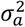 is the total additive genetic variance of the model.

### Models and analyses

#### Genome-wide associations

Efficient mixed-model association expedited (EMMAX) was performed using Gemma software [17], with the following model:

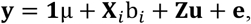

where **y** is a vector of phenotypes, μ is an overall mean, **X**_*i*_ is a vector of genotypes for *i*^*th*^ SNP, b_*i*_ is the substitution effect of the *i*^*th*^ SNP, **Z** is an incidence matrix for vector **u**, and **u** is a vector of random additive genetic effects, with 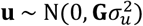, and **e** is a vector of residuals, with 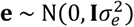 and **I** an identity matrix. The **G** was computed as in Zhou [17]:

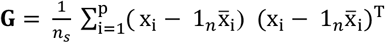

where the x_i_ is the *i*^*th*^ SNP locus column, 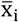 is the marker sample mean of the *i*^*th*^ locus, *n* and *n*_*s*_ are numbers of genotyped animals and SNP.

#### Genomic prediction

A linear mixed model was used to compute GP:

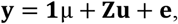

where **y** is a vector of phenotypes, μ is an overall mean, **Z** is an incidence matrix for vector **u**, and **u** is a vector of random additive genetic effects, with 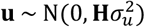 and **H** is the realized relationship matrix, and **e** is a vector of residuals, with 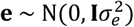. The GP was carried out with ssGBLUP and weighted ssGBLUP (WssGBLUP) [18] using the BLUPF90 family of programs [15]. For the mixed model equations in ssGBLUP and WssGBLUP, **H**^−1^ combines both pedigree and genomic relationships [19]:

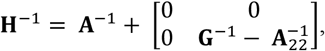

where **G**^−1^ is the inverse of the genomic relationship matrix and 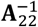 is the inverse of the pedigree relationship matrix for the genotyped animals. The **G** was created as in VanRaden [20]:

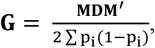

where **M** was defined before, p_i_ is the minor allele frequency of the *i*^*th*^ SNP, **D** is the diagonal matrix of SNP weights with dimensions equivalent to the number of SNP. In ssGBLUP, all SNP were assumed to have homogeneous weights, meaning that **D** was an identity matrix. To avoid singularity issues, **G** was blended with 5% of **A**_22_.

For the WssGBLUP, SNP effects were back-solved from GEBV (**û**) as described in Wang et al. [18]:

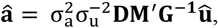

where **â** is a vector of estimated SNP effects, 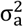 is the SNP variance, 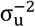 is the genetic variance, **D** is the diagonal matrix of SNP weights (**I** in ssGBLUP), and **M** is the centered matrix of genotypes. After SNP effects were estimated, the variance for the *i*^*th*^ SNP was calculated using the non-linearA method [20]:

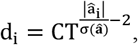

where CT is a constant that determines the departure from normality when deviating from 1; |â_i_| is the absolute estimated SNP effect for i^th^ SNP; and σ(**â**) is the standard deviation of the vector of estimated SNP effects. This study used CT as 1.125, an empirical value based on polygenic traits in dairy cattle populations. Results from the second iteration of weights were used in this study to maximize the prediction accuracy, as suggested by Zhang et al. [21], Lourenco et al. [22].

#### Validation of genomic predictions

In each scenario, prediction accuracy was calculated as the correlation between TBV and genomic estimated breeding value (GEBV). Besides, the regression coefficient (b_1_) of TBV on GEBV was used as an indicator of inflation or deflation of GEBV. When b_1_ is lower than one, it is indicative of inflation and deflation otherwise. As replicates were not used in this study, standard errors (SE) were computed using the bootstrapping method [23].

## Results

### Variant identification

The preliminary analysis showed similar results for both GWA and GP when QTN were included (withQTN) or not included in the data. Therefore, results of withQTN are only described. The results of GWA are shown in Fig. 1 ∼ 4. As most of the quantitative traits are highly polygenic, only results of Q2000 with H30 and H99 are shown with two *N*_*e*_ scenarios (*N*_*e*_20, *N*_*e*_200), respectively. In addition, the GWA results of EIG60, EIG70, and EIG80 were not included in those figures due to their insignificance. All other results with H90, Q200, and EIG60 ∼ EIG80 are described in the Additional file 1: Fig. 1. The number of significantly identified QTN, SNP, and the variance explained by those QTN of all scenarios above is in Additional file 1: Table 1. Considering a population with *N*_*e*_ equal to 20 and using EIG50, EIG90, EIG95, EIG98, and EIG99 as the sample size for GWA in Q2000 was not enough to detect significant QTN when the amount of information for genotyped animals was small (Fig. 1). However, when the sample size increased to 30,000 (i.e., ALL), three significant QTN were identified. In contrast, using genotyped animals with high reliability increased the ability to identify simulated QTN correctly (Fig. 2). EIG95 could capture three significant QTN, and as sample size increased to EIG98, EIG99, and ALL, 17, 33, and 142 QTN were identified, respectively.

**Figure 1.**
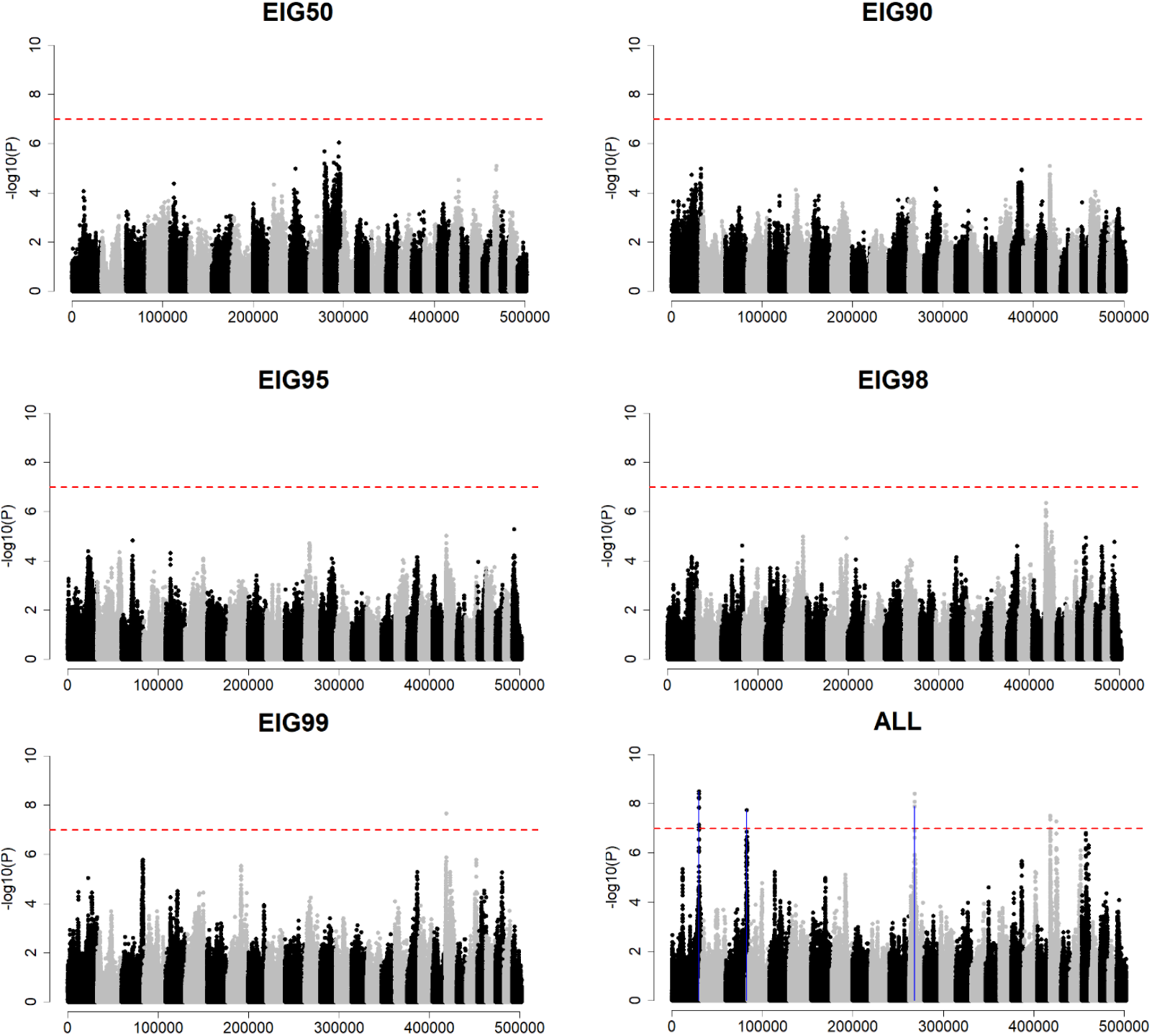
GWA results – EIG50, EIG90, EIG95, EIG98, EIG99, All – Ne20 Q2000 H30. *The x-axis and y-axis are indicating the number of variants and -log10(p-value). The red horizontal dot line represents the Bonferroni correction threshold. Blue vertical lines are pointing out the QTN position which was significantly identified.

**Figure 2.**
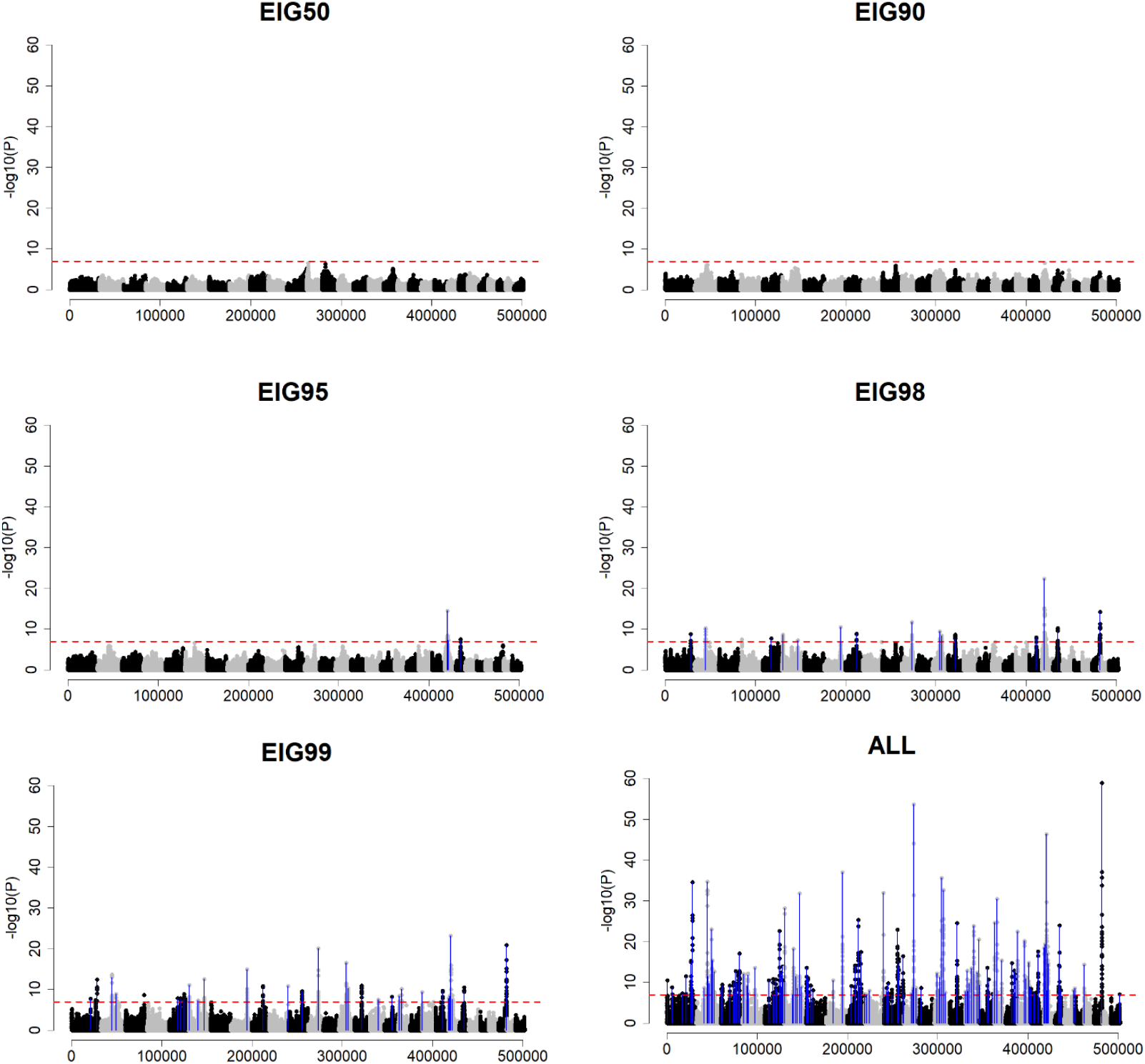
GWA results - EIG50, EIG90, EIG95, EIG98, EIG99, All – Ne20 Q2000 H99. *The x-axis and y-axis are indicating the number of variants and -log10(p-value). The red horizontal dot line represents the Bonferroni correction threshold. Blue vertical lines are pointing out the QTN position which was significantly identified.

Different patterns were observed for a population with *N*_*e*_ equal to 200 when having contrasting EIG*x* as the sample size (Fig. 3 and Fig. 4). Although EIG50, EIG90, and EIG95 were not sufficient to capture the significant QTN in H30, a sample size of EIG98 allowed the identification of seven QTN (Fig. 3). Moreover, increasing the number of genotyped animals to EIG99 and ALL helped detect more QTN and improve the GWA resolution, even though genotyped animals had low reliability. When *N*_*e*_ was 200, but the animals had high reliability, EIG90 is an adequate sample size to detect the QTN with the largest effect size (Fig. 4). For this scenario, EIG98 provided a clear resolution, similar to EIG99 and ALL. It is important to note that the number of largest eigenvalues explaining a certain proportion of the variance in **G** was different for *N*_*e*_20 and *N*_*e*_200 (Table 2). With all available genotyped animals (i.e., ALL), *N*_*e*_200 had more significant QTN discovered in GWA than *N*_*e*_20. For example, in Q2000 and H30, the three significant QTN in *N*_*e*_20 captured 3.9% of the additive genetic variance, whereas, in *N*_*e*_200, 15 QTN were capturing 13.9%. In both Ne scenarios, using genotyped animals with high reliability helped better detect QTN. For a less polygenic trait (Q200), fewer genotyped animals were required to identify the simulated QTN than in a more polygenic trait (Q200).

**Figure 3.**
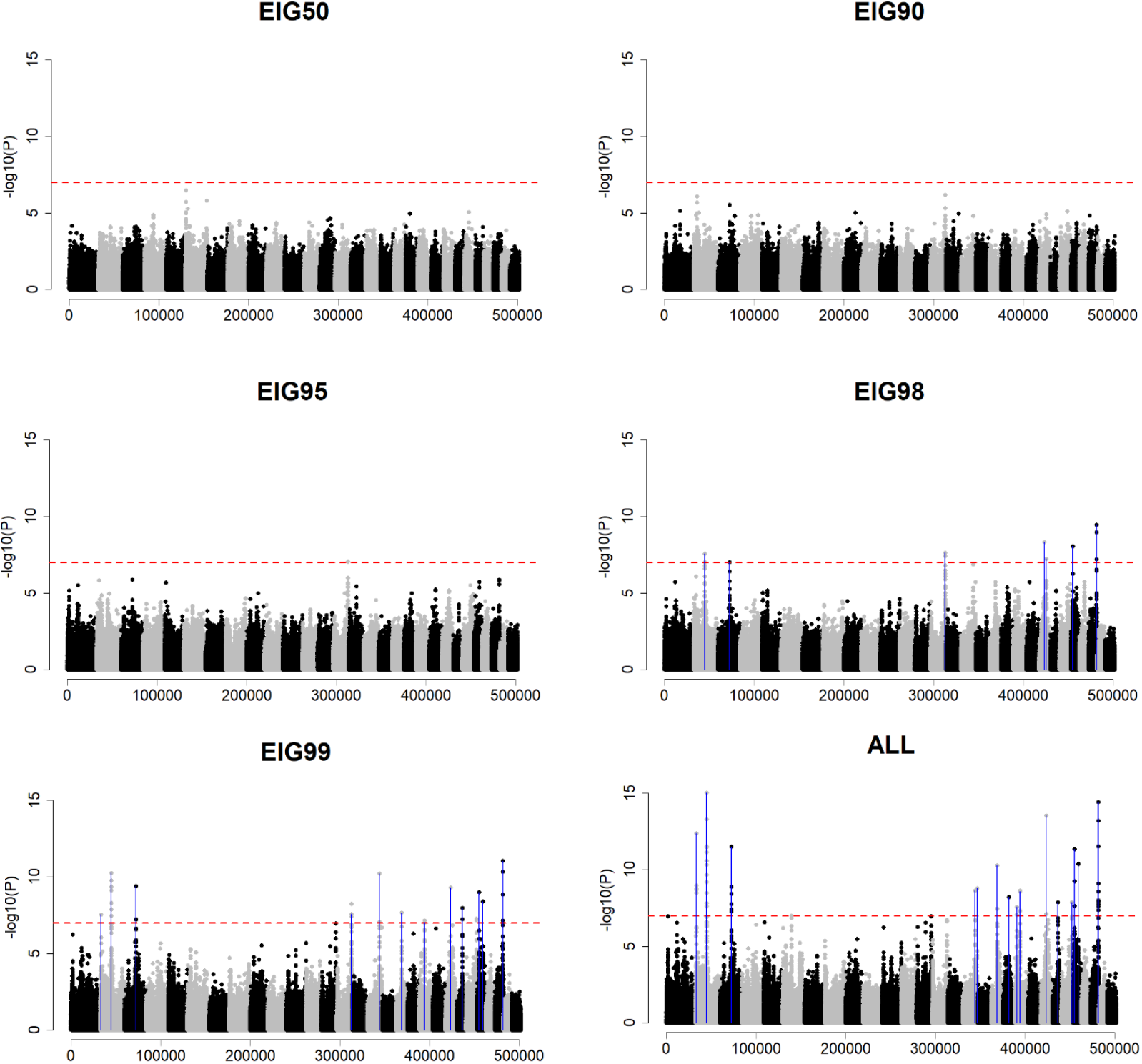
GWA results - EIG50, EIG90, EIG95, EIG98, EIG99, All – Ne200 Q2000 H30. *The x-axis and y-axis are indicating the number of variants and -log10(p-value). The red horizontal dot line represents the Bonferroni correction threshold. Blue vertical lines are pointing out the QTN position which was significantly identified.

**Figure 4.**
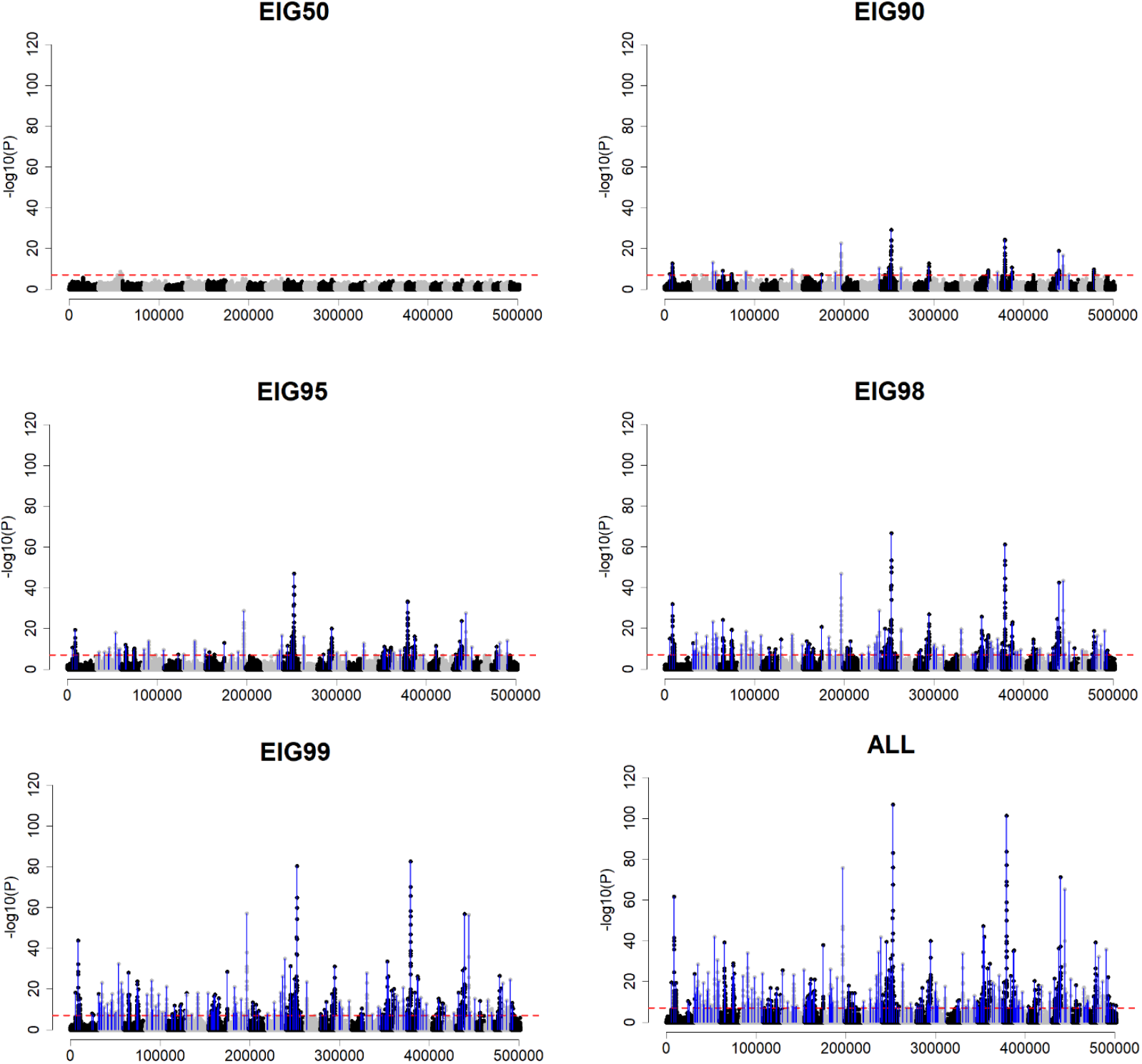
GWA results - EIG50, EIG90, EIG95, EIG98, EIG99, All – Ne200 Q2000 H99. *The x-axis and y-axis are indicating the number of variants and -log10(p-value). The red horizontal dot line represents the Bonferroni correction threshold. Blue vertical lines are pointing out the QTN position which was significantly identified.

### Association between variance explained by identified QTN and sample size

The proportion of variance explained by identified QTN is shown in Fig. 5. When the sample size increased, the proportion of variance explained by the identified QTN increased regardless of heritability, *N*_*e*_, and number of QTN. Comparing the proportion of variance explained according to the heritability, high heritability scenarios better identified significant QTN. For example, using all genotyped animals in *N*_*e*_20 and Q200 with H30 helped identify QTN explaining 44.9% of the variance (Fig. 5a). However, H90 and H99 identified QTN explaining 72.9% and 74.3% of the variance, respectively, with the same number of genotyped animals. For a more polygenic trait (Fig. 5b), QTN explained 3.9% and 43.1% of the variance with H30 and H99, respectively. The latter was similar to the variance explained in Q200, H30. Similar patterns were observed between *N*_*e*_20 and *N*_*e*_200; however, QTN identified in *N*_*e*_200 explained a greater proportion of variance than *N*_*e*_20 for Q200 and Q2000 (Fig. 5c-d). For example, the maximum proportion of variance explained by QTN in *N*_*e*_20 (Fig. 5a-b) was 74.3% and 43.1% for Q200 and Q2000 with H99, whereas those values were 96% and 65.1% in *N*_*e*_200 (Fig. 5c-d). One remarkable discovery with a less polygenic trait was that the use of EIG98 and EIG99 showed a similar proportion of variance explained by identified QTN as in ALL (Fig. 5a and 5c). For H99, when EIG98, EIG99, and ALL were used, the variance explained by the identified QTN was 59.9%, 68%, and 74.3%, respectively (Fig. 5a). In the case of Q2000, the scenarios mentioned above identified QTN explaining 11.2%, 18.9%, and 43.1% of the variance (Fig. 5b); therefore, the proportion of variance explained increased by almost fourfold from using EIG98 to ALL, whereas this increase was only 25% with Q200. In a larger population (*N*_*e*_200) and H99 (Fig. 5c), EIG98, EIG99, and ALL schemes detected QTN explaining 95.2%, 95.6%, and 96% of the variance for Q200. Even for the more polygenic scheme (Q2000), EIG98, EIG99, and ALL captured QTN explaining 52.5%, 59.7%, and 65.1% (Fig. 5d). Similar patterns were observed for the other two heritability scenarios in Fig. 5c-d.

**Figure 5.**
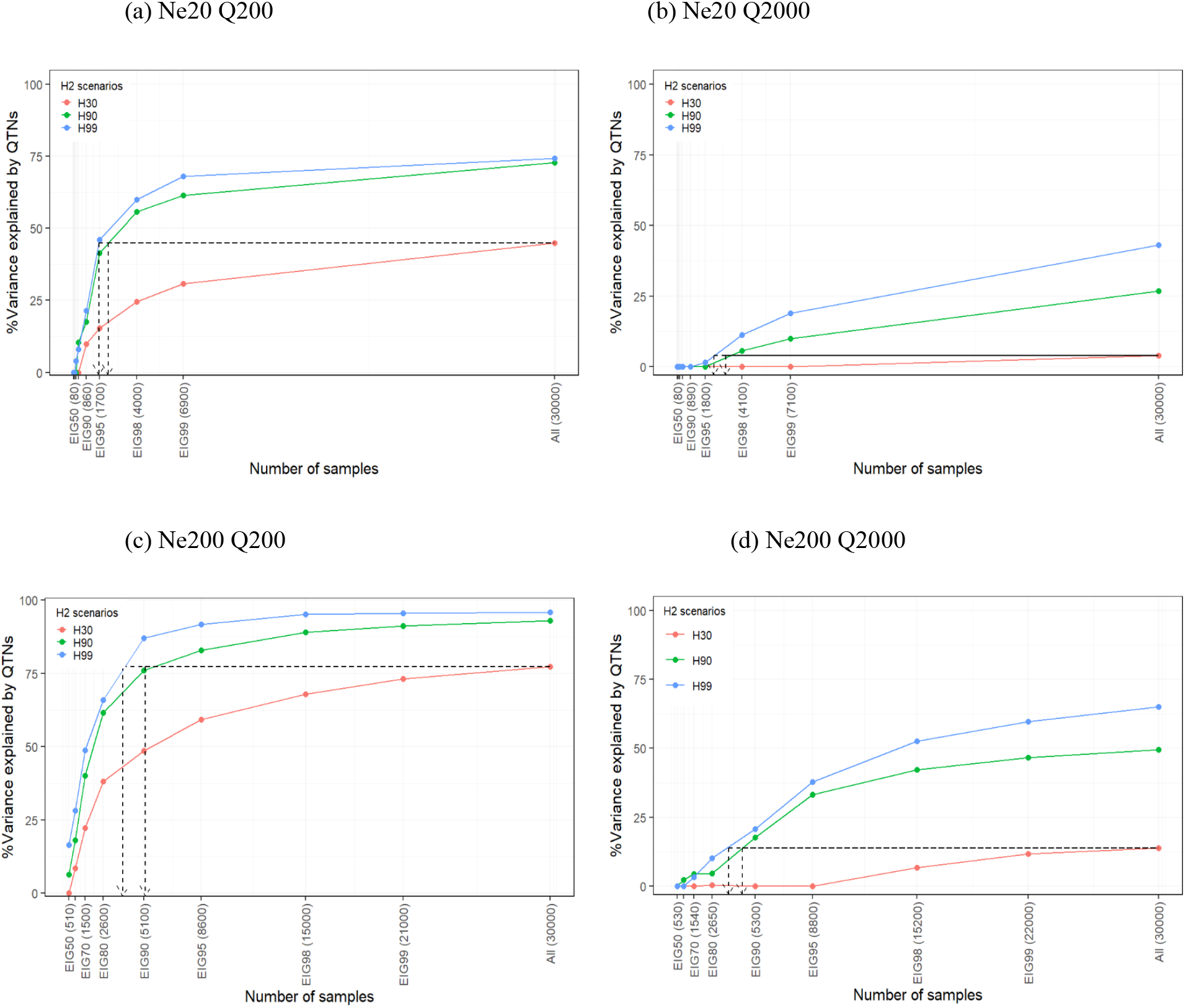
Total variance explained by significant QTN across the different sample sizes and heritabilities. *Each x-axis and y-axis indicate the number of genotyped animals (EIGx scenarios) and percentage of total variance explained by QTN for three heritability scenarios (H0.3, H0.9, and H0.99).

To investigate the corresponding sample size from the use of low to high heritability scenarios at the point where the same percentage of variance was explained, we used H30 as a benchmark. Table 3 shows the approximated sample size when the largest discovery set (ALL, N = 30,000) was used. All approximated sample sizes based on other discovery sets as a benchmark are in Additional file 1: Table 2. The results of the *N*_*e*_20 Q200 scenario showed that around 2223 and 1662 genotyped animals were required for H90 and H99 to reach the same magnitude of genetic variance explained by the identified QTN in H30 (Table 3). In *N*_*e*_20 Q2000, using 3049 and 2378 genotyped animals with H90 and H99 helped identify QTN explaining 3.9% of the variance, which was accomplished using ALL in H30. Similar patterns were observed in *N*_*e*_200 scenarios. One remarkable difference between *N*_*e*_20 and *N*_*e*_200 was that the sample size to reach the same proportion of variance explained by QTN using ALL in H30 was equivalent to EIG90 ∼ EIG98 in *N*_*e*_20 but EIG80 ∼ EIG90 in *N*_*e*_200 when considering H90 and H99.

**Table 3.** Approximated sample size based on local polynomial regression and proposed equation using ‘ALL’ as benchmark.

| Scenario | Heritability | %Var<sup>1</sup> | SS<sub>pol</sub><sup>2</sup> | EIGx<sub>app1</sub><sup>3</sup> | SS<sub>rel</sub><sup>4</sup> | EIGx<sub>app2</sub><sup>5</sup> | Diff<sup>6</sup> |
| --- | --- | --- | --- | --- | --- | --- | --- |
| Ne20 Q200 | 0.3 | 44.9 | 30000 |  |  |  |  |
|  | 0.9 | 44.9 | 2223 | EIG95~98<br>(1770~4100) | 2698 | EIG95~98<br>(1770~4100) | 475 |
|  | 0.99 | 44.9 | 1662 | EIG90~95<br>(890~1800) | 2453 | EIG95~98<br>(1800~4100) | 791 |
| Ne20 Q2000 | 0.3 | 3.9 | 30000 |  |  |  |  |
|  | 0.9 | 3.9 | 3049 | EIG95~98<br>(1750~4100) | 3007 | EIG95~98<br>(1750~4100) | -42 |
|  | 0.99 | 3.9 | 2378 | EIG95~98<br>(1700~4000) | 2734 | EIG95~98<br>(1700~4000) | 356 |
| Ne200 Q200 | 0.3 | 77.4 | 30000 |  |  |  |  |
|  | 0.9 | 77.4 | 5205 | EIG80~90<br>(2650~5250) | 4324 | EIG80~90<br>(2650~5250) | -881 |
|  | 0.99 | 77.4 | 3814 | EIG80~90<br>(2700~5300) | 3931 | EIG80~90<br>(2700~5300) | 117 |
| Ne200 Q2000 | 0.3 | 13.9 | 30000 |  |  |  |  |
|  | 0.9 | 13.9 | 4524 | EIG80~90<br>(2600~5100) | 4719 | EIG80~90<br>(2600~5100) | 195 |
|  | 0.99 | 13.9 | 3664 | EIG80~90<br>(2600~5100) | 4290 | EIG80~90<br>(2600~5100) | 626 |

\*%Var1: percentage of variance explained by significantly identified QTN
\*SS<sub>pol</sub>2: Approximated sample size using local polynomial regression
\*EIGx<sub>app1</sub>3: EIGx scenario range including Sample<sub>app1</sub>
\*SS<sub>rel</sub>4: Approximated sample size using proposed equation
\*EIGx<sub>app2</sub>5: EIGx scenario range including Sample<sub>app2</sub>
\*Diff6: Difference between Sample<sub>app2</sub> and Sample<sub>app1</sub>
\*Dotted arrows in Fig. 2 matched with each line of H90 and H99 scenario was approximated sample size using local polynomial regression and those were described above as Sample<sub>app1</sub>.
\*EIGx scenario ranges including either Sample<sub>app1</sub> (EIGx<sub>app1</sub>) or Sample<sub>app2</sub> (EIGx<sub>app2</sub>) are described with the difference between Sample<sub>app1</sub> and Sample<sub>app2</sub> as Sample<sub>app2</sub> – Sample<sub>app1</sub> (Diff) in Table

In the current study, in addition to the approximated sample size by local polynomial regression, we derived the equation to estimate the sample size relating the number of samples, *M*_*e*_, *N*_*e*_, percentage of variance explained by the identified QTN, and reliability of EBV. For example, to approximate the sample size of H90 in *N*_*e*_20 Q200 scenario when %var was 44.9, it would be

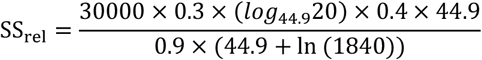

Those sample sizes approximated by the proposed equation are described in Table 3. Approximating the sample size in Q2000 was more accurate than in Q200 for *N*_*e*_20 and *N*_*e*_200. Additionally, the formula and the polynomial regression provided sample sizes that were within the same EIGx range, except for one scenario (*N*_*e*_20 Q200 H99). As an example, *N*_*e*_20 Q2000 resulted in 3049 (H90) and 2378 (H99) samples in SS_pol_ and 3007 (H90) and 2734 (H99) in SS_rel_; however, the sample size in both cases laid within EIG95∼98. Differences between SS_pol_ and SS_rel_ across more polygenic scenarios were not large.

### Genomic predictions

Beforehand, we investigate the possible bias of GP by using the same set of genotyped animals for both discovery and training. Using different groups of genotyped animals for discovery and training resulted in less inflation of GEBV than utilizing the same animals for both processes (results are not shown). Therefore, the GP analyses were done with training animals different from the discovery set.

We noticed very small standard errors for prediction accuracy (<0.005) and b_1_ (<0.01) by bootstrapping. The results of prediction accuracy and inflation/deflation indicator of GEBV (b_1_) are shown in Fig. 6 and Fig. 7, respectively. Those accuracies and b_1_ were calculated as the average of all genotyped scenarios: 50k, TOP10, TOP50, TOP100, TOP200, TOP400, and ‘SIG’ for Q200 and 50k, TOP10, TOP100, TOP500, TOP1000, TOP2000, TOP4000, and ‘SIG’ for Q2000 scenarios. Standard deviations were less than 0.02 for all scenarios. Fig. 6 shows the prediction accuracy depending on the number of QTN (Q2000 or Q200), trait heritability (H30, H90, and H99), and training data scenarios (EIGx and ALL). In general, as the training set size increased, prediction accuracy also increased. Different patterns were observed between *N*_*e*_20 and *N*_*e*_200. In *N*_*e*_200, the prediction accuracy increased consistently as the training set size increased; however, the increase in prediction accuracy was not constant in *N*_*e*_20. For example, in *N*_*e*_20 Q200 H30, while the training data set went up from the EIG50 to EIG95, prediction accuracy increased only by about 0.03. A similar pattern was observed for *N*_*e*_20 Q2000 H30, which showed a gain of about 0.02. In general, *N*_*e*_20 showed greater prediction accuracy than *N*_*e*_200. For example, when the smallest sample size (EIG50) was used, the average accuracy in *N*_*e*_20 was 0.82±0.03 and *N*_*e*_200 was 0.74±0.09, a difference of about 0.08. This difference became smaller with the largest sample size (ALL), which was 0.04. Therefore, no substantial differences in prediction accuracy between the *N*_*e*_20 and *N*_*e*_200 were found when the largest training set was used. Overall, the difference in prediction accuracy between EIG98 or EIG99 and ALL was smaller in *N*_*e*_200 (0.88, 0.90, and 0.92 on average for EIG98, EIG99, and ALL) than in *N*_*e*_20 (0.90, 0.92, and 0.96 on average for EIG98, EIG99, and ALL), indicating a sample for training with the size of EIG98 or EIG99 would suffice.

**Figure 6.**
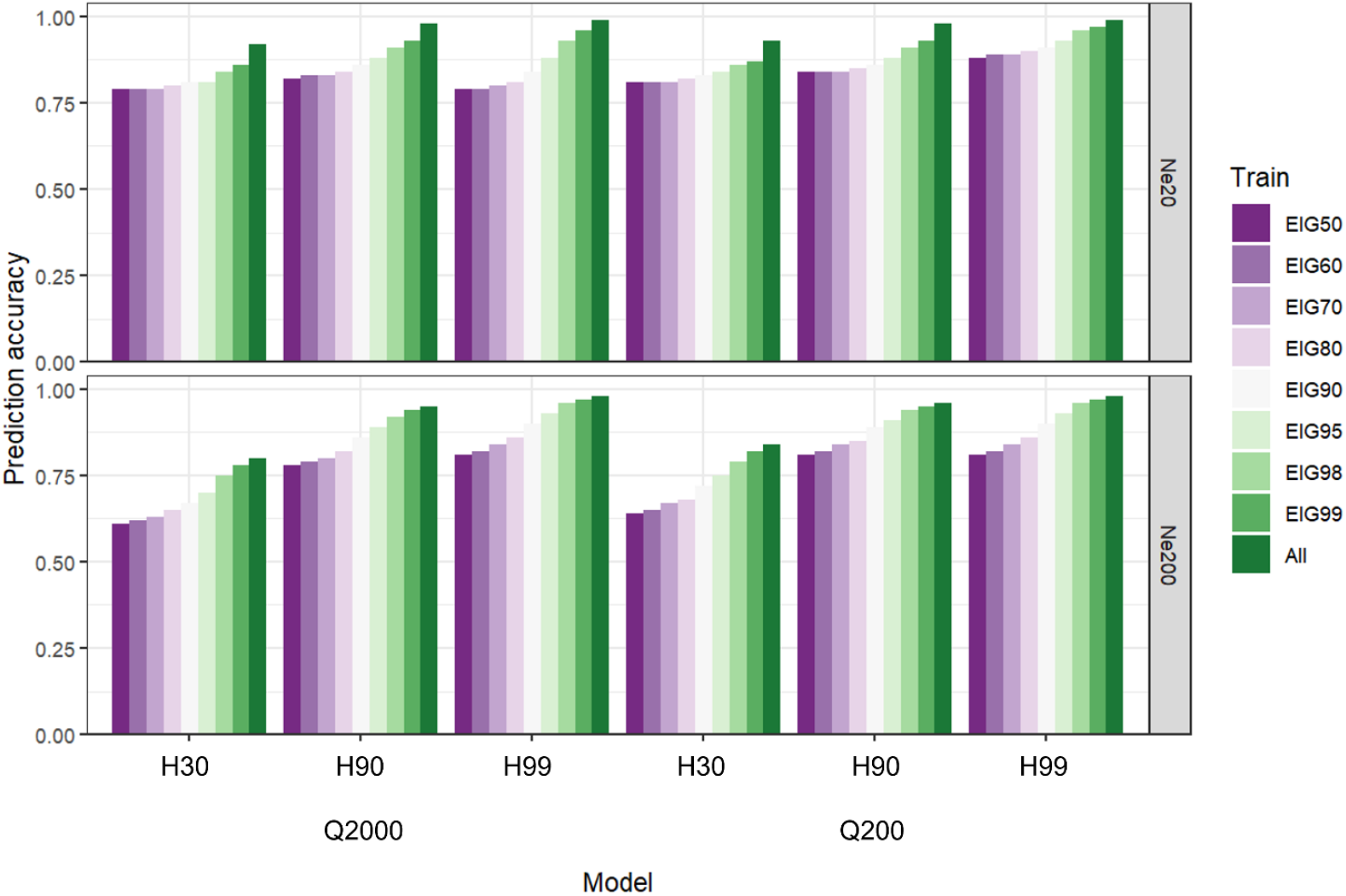
Prediction accuracy.

**Figure 7.**
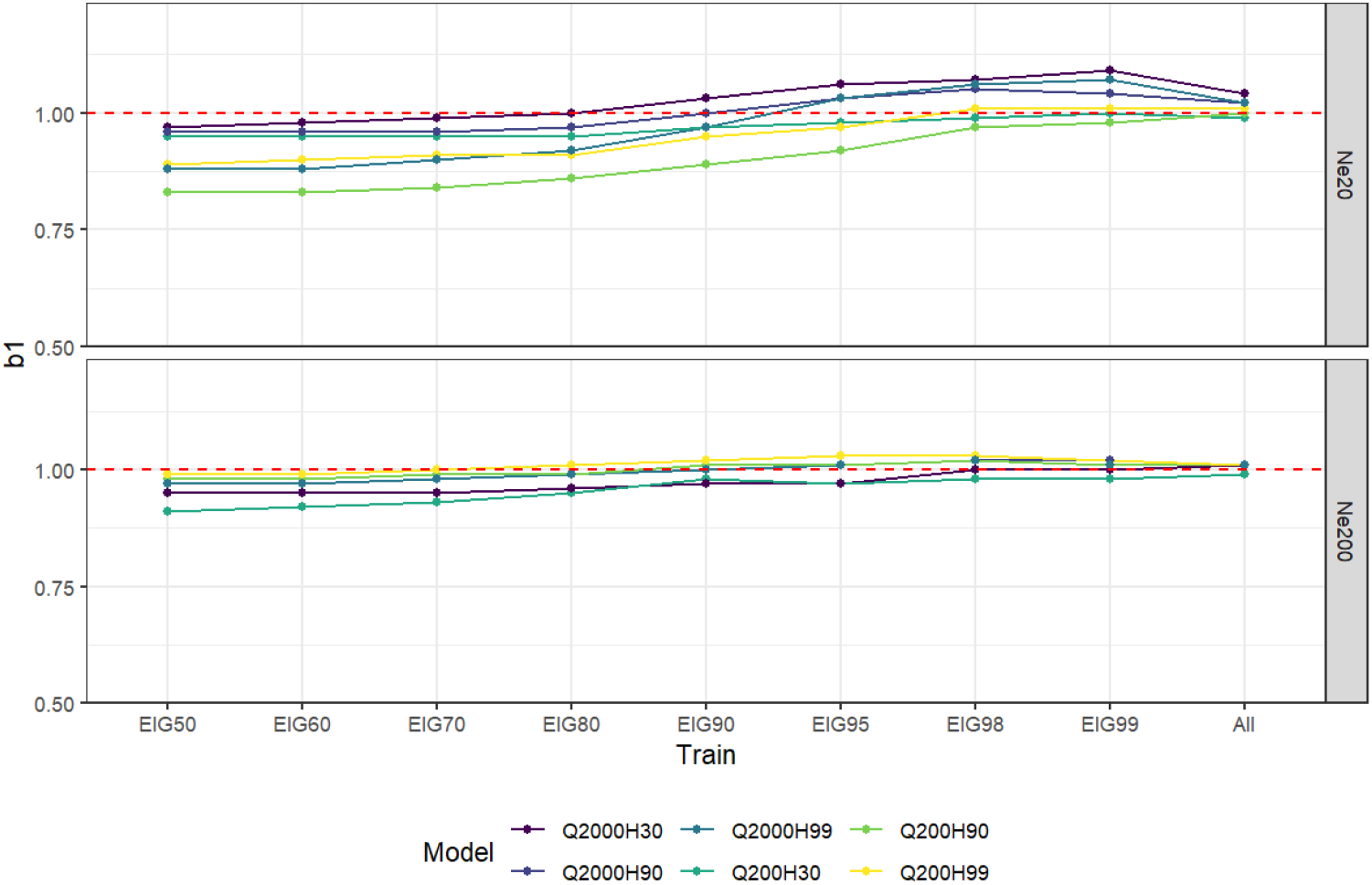
Regression coefficients (b1)

The number of QTN marginally affected the prediction accuracy for *N*_*e*_20 and *N*_*e*_200 scenarios. For both *N*_*e*_ scenarios, Q200 showed greater prediction accuracy, especially for the low heritability with the smaller training sets. For example, in *N*_*e*_20 H30, the prediction accuracy from EIG50 to EIG70 was 0.81 with Q200 and 0.79 with Q2000. With larger training set sizes and higher heritability, the difference between Q200 and Q2000 decreased. Prediction accuracies were highly influenced by the heritability of the trait, particularly with larger effective population size. For instance, with *N*_*e*_200 Q200, when EIG50 was used, the prediction accuracy was 0.64, 0.81, and 0.81 for H30, H90, and H99, respectively. Even with the most extensive training set (ALL), prediction accuracy was 0.84, 0.96, and 0.98, following the previous order. Similar patterns were observed for *N*_*e*_200 Q2000. Thus, using low to high-reliability genotyped animals for training could affect the prediction accuracy more in populations with large *N*_*e*_. Expanding the training set from EIG95 to EIG98 in *N*_*e*_200 H30 increased prediction accuracy by 5.5% and 6.82% for Q200 and Q2000, respectively. However, the maximum increase for *N*_*e*_200 H90 and H99 was observed when the training set size expanded from EIG80 to EIG90 (3.8% ∼ 4.61%). Unlike *N*_*e*_200, great improvements in accuracy were observed when moving from EIG99 to ALL in H30 and H90 for both Q200 and Q2000.

Regression coefficients (b_1_) used as indicators of inflation/deflation of GEBV are shown in Fig. 7. When the training set size was small, less inflation was observed in *N*_*e*_200 than in *N*_*e*_20. In both *N*_*e*_ scenarios, using a large training set alleviated the inflation, so when ALL was used for training, all scenarios had b_1_ close to 1, especially for *N*_*e*_20. Interestingly, more variation between the models was observed in *N*_*e*_20 than in *N*_*e*_200.

Fig. 8 shows the prediction accuracy with 50k compared to 50k plus SIG, TOP400 (Q200), and TOP4000 (Q2000), together with the percentage of gain by adding possible causative variants. As the increase was not major across analyses combining the 50k and the top SNP, only the scenarios with the largest changes are shown in Fig. 8; all the other scenarios are in Additional file 2. Overall, the percentage of gain was greater in *N*_*e*_200 (3.29% ∼ 9.01%) than in *N*_*e*_20 (0.86% ∼ 1.98%). In addition, Q200 showed a higher percentage of gain than Q2000 in both *N*_*e*_ scenarios. Interestingly, the maximum accuracy gain was usually observed when the largest number of top SNP (TOP400 for Q200 and TOP4000 for Q2000) was added to 50k chip data, representing twice the number of simulated QTN. The only exceptions were Q200 H90 and H99 for *N*_*e*_20 and *N*_*e*_200, which had the highest accuracy gain with 50k plus SIG. This is probably because identifying significant QTN was easier in a less polygenic trait (Q200) with H90 and H99. In contrast, finding QTN in Q2000 or a low heritability trait was harder.

**Figure 8.**
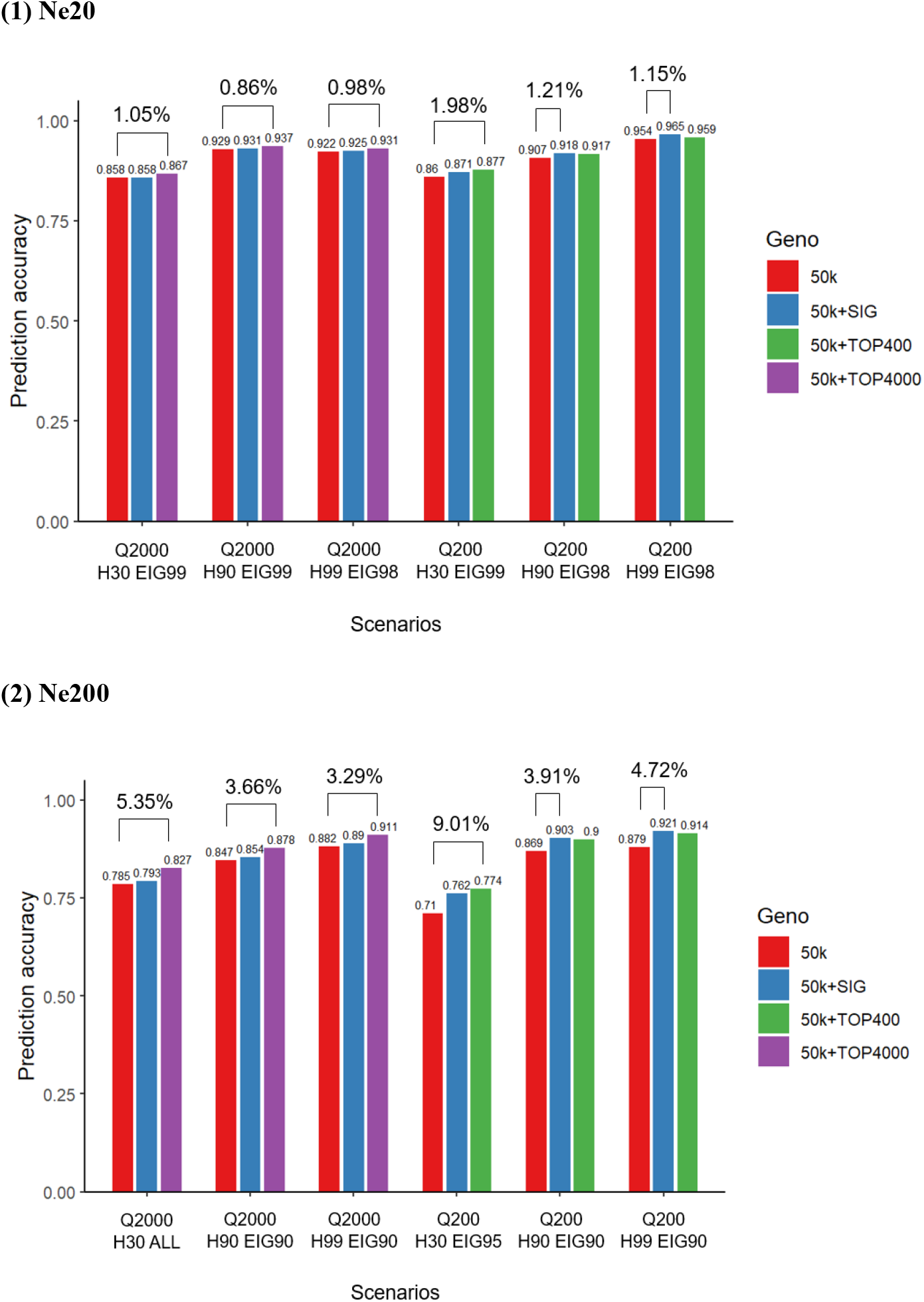
Prediction accuracy of 50k and TOPv scenario which showed maximum gain (%)

Assuming we know all the actual QTN, using them for GP returned the best predictive accuracy (Additional file 1: Fig. 2). However, it is hard to identify the real QTN using real data; therefore, the current study compared the prediction accuracy using TOP200 and TOP2000, QTN200 and QTN2000, and added those variants to the 50k panel to investigate the losses from having variants selected based on GWA. This was done for H99 when the selected variants were from using ALL (30,000 animals) as discovery and training sets. When only TOP200 (2000) and QTN200 (2000) were used, the difference in prediction accuracies was quite large (0.05 ∼ 0.07). However, prediction accuracies were almost the same when each TOP200 (2000) and QTN200 (2000) were combined with the 50k chip. Based on our findings, if QTN are in the data, using the top variants from GWA in GP can provide reasonable accuracy.

## Discussion

In this study, we comprehensively investigated the impact of using different sample sizes in GWA based on the dimensionality of the genomic information, the implications of using genotyped animals having low to high reliability of EBV in GWA, and the inclusion of preselected variants into a typical 50k SNP panel using ssGBLUP and WssGBLUP. These investigations brought insights into how different data structures can affect the performance of GWA and GP under the ssGBLUP framework. We used the concept of limited dimensionality of the genomic information [4]. Our results showed that this concept could be a helpful indicator of the number of genotyped animals required for GWA, depending on *N*_*e*_, *M*_*e*_, the number of QTN, and the reliability of EBV. Our results showed that having a sample size with the number of genotyped animals corresponding to that of EIG98 was appropriate for variant discovery, particularly in the population with large *N*_*e*_. Additionally, using genotyped animals with high EBV reliability could help better identify significant QTN regardless of *N*_*e*_ and the genetic architecture of the traits. Incorporating selected variants obtained from GWA to the 50k SNP chip could improve prediction accuracy when a training set with proper size was used; however, the gain could be limited in some scenarios.

### GWA – preselection of variants

The most prevalent workflow for GP with sequence data is 1) pre-selection of significant variants, 2) incorporation of selected variants to the commercial chip data (i.e., 50k) or fitting separate genomic matrices in the model [9, 24, 25], 3) comparison of the GP performance with a benchmark SNP chip. Several studies have been conducted to improve GP using sequence data with either simulated or real data. However, conclusions about the advantage of using sequence data have not been very consistent in the literature, and they seem to be dependent on several factors such as the species, the genetic architecture of the trait, the size of data, and statistical methods [9, 24, 26, 27].

Among those factors, the most critical is the size of data for discovery, training, and test sets. Specifically, the sample size for the variant discovery set is essential as it is the first step and, thus, predominantly affects the results of the entire study. Current results indicated that using a small number of genotyped animals could not identify the significant SNP or QTN. Lourenco et al. [22] used two different numbers of genotyped animals (N = 2,000 and 25,000) for GWA and reported that the best resolution was observed when more genotyped animals were used. In the same line, de Las Heras-Saldana et al. [28] outlined that using a larger dataset for GWA allowed to better identify quantitative trait loci (QTL) regions for carcass traits in Hanwoo cattle.

As the number of genotyped animals has currently increased in many species, for instance, about 5 million U.S. Holsteins (https://queries.uscdcb.com/Genotype/cur_freq.html), and about 1 million American Angus (K. Retallick, American Angus Association, Saint Joseph, MO, personal communication) have been genotyped as of March 2022, it is important to know how many genotyped animals are effectively required to detect the significant variants. Current results showed that using at least the number of genotyped animals equivalent to EIG98 could identify the most informative QTN. Using EIG99 or all available genotyped animals little improved the ability to identify significant QTN in *N*_*e*_200 for both Q200 and Q2000 scenarios. This result could be helpful for both small and large genotyped populations with large *N*_*e*_. For breeding populations with fewer resources, the number of animals to genotype may be limited; therefore, accessing what would be the effective sample size could benefit cost-effective genotyping or sequencing. For large populations, our study showed that not all animals are needed for variant discovery, and a balanced data set should be constructed for discovery, training, and testing to avoid biases and maximize the power to detect the significant variants. When *N*_*e*_ was small and the trait was highly polygenic, the small sample size could not identify any significant QTN until it reached ALL and EIG98 for H30 and H90. With more information on genotyped animals (i.e., H99), a sample size equivalent to EIG95 helped identify a few QTN. Chicken and pigs had smaller *N*_*e*_ (32 ∼ 48) than cattle among the livestock species [29]. Therefore, using a sample size corresponding to less than ALL would not be enough to detect significant signals for those species. Gozalo-Marcilla et al. [30] carried out large-scale GWA for backfat thickness in pigs using around 15k to 55k genotyped animals. They found 264 significant SNP across 8 different lines for traits with moderate to high heritability (0.30 ∼ 0.58). As backfat thickness has been known for its polygenic architecture (more than 1400 QTL associated backfat thickness is reported in https://www.animalgenome.org/QTLdb), their discovery is supported by our findings in populations with small *N*_*e*_ and moderate heritability.

Pocrnic et al. [4] described the number of largest eigenvalues explaining a certain proportion of **G** as a function of *N*_*e*_ and genome length in Morgans, such that EIG90 ≈ *N*_*e*_*L*, EIG95 ≈ 2*N*_*e*_*L*, and EIG98 ≈ 4*N*_*e*_*L*. Stam [3] expressed the expected number of independent chromosome segments as 4*N*_*e*_*L*. In this study, *N*_*e*_ was 20 or 200 and *L* was 23, so *M*_*e*_ was approximated as 1,840 and 18,400, between EIG95 and EIG98 for *N*_*e*_20, and EIG98 and EIG99 for *N*_*e*_200 in Table 2. As *N*_*e*_ and *M*_*e*_ are proportional, smaller *N*_*e*_ denotes fewer *M*_*e*_, indicating fewer blocks are existed in the genome with a strong LD between variants because of the close relationship among individuals. *N*_*e*_ also has been reported as a factor affecting the performance of GWA [22, 31]. In the current study, we showed that when the same number of genotyped animals were used (ALL), *N*_*e*_200 could better identify the significant QTN explaining more genetic variance than *N*_*e*_20 for all heritability and QTN scenarios. This might be because of smaller chromosome segments and weaker LD between the QTN and SNP in *N*_*e*_200 than in *N*_*e*_20. Pinpointing QTN is harder in *N*_*e*_20 because many SNP may capture the QTN signal. The noise in GWA resolution is possibly due to the strong relationships between the SNP and QTN, established by a highly structured population over the generations in *N*_*e*_20; therefore, identifying the true causative variant is not trivial in smaller populations. In general, *N*_*e*_ of farm animals such as chickens, pigs, dairy, and beef cattle is less than 200 and could range from 40 to 150 [29]; thus, the current findings would be helpful information for the future GWA in those species. However, identifying all significant variants is not assured due to the polygenic nature of most traits in livestock animals, and most of the causal variants have a small effect. For example, even with the largest number of genotyped animals for GWA in our study (ALL, N = 30k), identifying QTN with very small effects was not possible due to limited statistical power. Misztal et al. [12] showed that identifying all simulated QTN was impossible even when all had the same effect, *N*_*e*_ was 600, and the sample size was 6000. For a population with *N*_*e*_ of 60, a sample size three times larger resulted in more true signals in GWA, but not as many as with *N*_*e*_ of 600. The same authors argued that with smaller *N*_*e*_, more data is required to overcome the noise stage and capture the actual signals.

Different heritability scenarios were compared to investigate the performance of GWA when genotyped animals with low to high reliability of EBV were used for the variant discovery stage. Our findings highlighted that regardless of the number of QTN, *N*_*e*_, *M*_*e*_, and sample size, high heritability scenarios could capture more significant QTN explaining a larger portion of the variance. However, Takeda et al. [32] observed no differences in power to detect QTL when heritability 0.2 and 0.5 were simulated but outlined that QTL detection was better with the increasing number of phenotyped progenies (N = 1,500, 4,500, 9,000). As the use of more phenotyped progeny data indicated higher reliability of EBV of the parents, which is the case of genotyped animals in the higher heritability scenarios in our study, those findings agreed with the current results. Besides, van den Berg et al. [33] reported that the number of false positives in QTL detection decreased with increasing heritability and number of records. Thus, using genotyped animals with high EBV reliability could sufficiently detect the QTN although few animals were used.

#### Sample size approximation

Approximated sample sizes were obtained through polynomial regression and a formula relating *N*_*e*_, *M*_*e*_, and percentage of variance to be explained by significant QTN. The latter was useful to investigate the sample size given the average reliability of EBV in the set of animals available for GWA. Overall, the comparison showed a better approximation with Q2000 scenarios but a very different approximation scale with Q200 scenarios (Additional file 1: Table 2). One possible reason for the inaccurate approximated sample size would be an unbalanced simulation design for *N*_*e*_ and heritability. As we only simulated two scenarios of *N*_*e*_: 20 and 200 for the smallest and largest *N*_*e*_ in livestock species with three heritability scenarios: 0.3, 0.9, and 0.99 representing low, high, and very high reliability of EBV, there is a large gap between *N*_*e*_ 20 and *N*_*e*_200, and an irregular pattern of heritability scenarios. However, the proposed equation could be applied for more polygenic traits. For example, given the number of genotyped or sequenced animals available, reliability of those animals, reliability of target animals, and proportion of variance explained by identified QTN, i.e., identified SNP in real data, with *N*_*e*_ and *M*_*e*_, we can approximate the sample size for GWA.

#### Genomic prediction

In general, the accuracy of GP was improved as the size of training data increased, and combining selected variants to a 50k SNP panel could improve accuracy when the GP was performed with the proper size of training sets. It was demonstrated that increasing the number of animals in training sets improved the accuracy of GP [34-36]. Our findings support those results, although only a tiny improvement (< 1.0%) was reported when using training sets with the number of genotyped animals equal to EIG50 to EIG70 in *N*_*e*_20. Moser et al. [37] observed no improvements in prediction accuracy when the training size was enlarged from 1,239 to 1,880 in Australian dairy cattle. Therefore, adding a substantial number of genotyped animals to the training set is necessary to improve prediction accuracy. As the current study suggested, the training set size can be based on the number of eigenvalues explaining a certain percentage of the variance in **G**. Those improvement patterns were very similar in both *N*_*e*_20 and *N*_*e*_200; however, the prediction accuracies were generally smaller for the *N*_*e*_200 when the same number of genotyped animals were used. Daetwyler et al. [38] investigated the impact of the genomic structure of the population (*N*_*e*_ and *M*_*e*_) on the accuracy of GBLUP. In their study, with the same number of individuals in the training sets, smaller *N*_*e*_ showed better accuracy than larger *N*_*e*_ regardless of the number of QTL.

Daetwyler et al. [34] proposed the following equation for prediction accuracy: 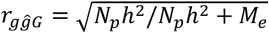, where *N*_*p*_ is a training set size, *h*^2^ is the trait heritability, and *M*_*e*_ is the number of independent chromosome segments. Equation to approximate the *M*_*e*_ proposed by Stam [3] was 4*N*_*e*_*L*, where *M*_*e*_ is proportional to *N*_*e*_, thus current results that showed greater accuracy with smaller *N*_*e*_ theoretically sounded when the *N*_*p*_ and *h*^2^ are equivalent. In addition, the small size of *N*_*e*_ means that fewer *M*_*e*_ to estimate, thus a smaller prediction error variance would be estimated [4].

We selected variants based on a p-value of 0.05 with a Bonferroni correction for multiple testing and the order of the significance level; however, the Bonferroni correction might generate a stringent threshold, increasing the number of false negatives. Therefore, we tested GP by combining selected variants based on sample size (TOP*v*) and significant variants. We demonstrated that when a large training set incorporated a relatively large number of variants (i.e., twice the number of simulated QTN), prediction accuracy improved by up to 9%. Several studies used selected variants from imputed sequence data to improve GP in single-breed populations. Veerkamp et al. [11] reported that when selected variants were used for GP, accuracy decreased, and bias increased. However, VanRaden et al. [26] observed an improvement in accuracy up to 5% when 16k selected variants were added to 60k chip data. In single-breed populations, an improvement in prediction accuracy using selected variants from sequence data could be limited due to long-range LD; thus, precise identification of variants is much harder than multi-breed or across-breed [11].

Fragomeni et al. [7] outlined that including causative QTN in the unweighted **G** through ssGBLUP increased accuracies by 0.04 when the number of QTN was 100 and 1000, which was similar to our results (0.02 ∼ 0.06). Additionally, when those authors added weights derived from SNP effects to **G**, accuracies increased by 0.10 and 0.03 for 100 and 1000 QTN scenarios, respectively, meaning that weighting SNP was more important for the scenario with a smaller number of QTN (less polygenic). However, in the current study, WssGBLUP resulted in no improvement in accuracy, and more inflation of GEBV was observed compared to ssGBLUP. However, the inflation of GEBV was reduced when more genotyped animals were added to the training set. The major difference between ssGBLUP and WssGBLUP is that ssGBLUP assumes that all SNP explain the same amount of genetic variance, whereas WssGBLUP assigns different variances for each SNP [18]. In general, weighting **G** may not increase the accuracy of GP but may improve the GWA resolution (Wang et al., 2012).

In our study, the resolution of variant detection was improved using at least the number of genotyped animals corresponding to the number of eigenvalues explaining 98% of the variation in **G** for the *N*_*e*_200 scenarios, meaning that when the number of genotyped animals for discovery is close to the approximated *M*_*e*_, precise detection of significant variants is feasible. As the genomic information has limited dimensionality, it could be expressed as the number of non-redundant SNP, genotyped animals, and *M*_*e*_ [39]. Therefore, investigating the dimensionality of the genomic information can help determine the sample size required for discovery and training. Since the performance of GWA and GP depends on several factors such as the genetic architecture of the trait, population structure, heritability, and sample size, more research is needed with real data to validate our results.

## Conclusion

Accurately identifying causative variants from sequence data depends on the effective population size and, therefore, the dimensionality of genomic information. This dimensionality can help identify the suitable sample size for GWA and should be considered for variant selection. Assigning genotyped animals with high breeding value reliability to the discovery set helps better identify the significant QTN. As sequence data become available, preselecting variants, and adding them to regular chip data could improve prediction accuracy if the dimensionality of the genomic information is considered; however, the improvement is mostly limited.

## List of abbreviations

A: pedigree relationship matrix
EBV: estimated breeding value
EIG: eigenvalues of **G**
EMMAX: efficient mixed-model association expedited
G: genomic relationship matrix
GEBV: genomic estimated breeding value
GP: genomic prediction
GWA: genome-wide association
H: realized relationship matrix
I: identity matrix
LD: linkage disequilibrium
Me: independent chromosome segments
Ne: effective population size
QTL: quantitative trait loci
QTN: quantitative trait nucleotides
ssGBLUP: single-step genomic best linear unbiased prediction
SE: standard errors
SNP: single nucleotide polymorphisms
TBV: true breeding value
WssGBLUP: weighted ssGBLUP

## Declarations

### Ethics approval and consent to participate

Not applicable.

### Consent for publication

Not applicable.

### Availability of data and materials

The datasets used and/or analysed during the current study are available from the corresponding author on reasonable request.

## Author’s information

### Affiliations

Department of Animal and Dairy Science, University of Georgia, Athens, GA, 30602, USA Sungbong Jang, Shogo Tsuruta, Natalia Galoro Leite, Ignacy Misztal & Daniela Lourenco

### Contributions

DL and SJ conceived and designed the study. SJ analyzed the data and wrote the first draft of the manuscript. DL helped with the computations and structure of the manuscript. NG, ST, IG, and DL provided critical insights and revised the manuscript. All authors read and approved the final manuscript.

## Literature Cited

1. Visscher PM, Wray NR, Zhang Q, Sklar P, McCarthy MI, Brown MA, et al. 10 years of GWAS discovery: biology, function, and translation. The American Journal of Human Genetics. 2017;101(1):5–22.

2. Berisa T, Pickrell JK. Approximately independent linkage disequilibrium blocks in human populations. Bioinformatics. 2016;32(2):283.

3. Stam P. The distribution of the fraction of the genome identical by descent in finite random mating populations. Genetics Research. 1980;35(2):131–55.

4. Pocrnic I, Lourenco DA, Masuda Y, Legarra A, Misztal I. The dimensionality of genomic information and its effect on genomic prediction. Genetics. 2016;203(1):573–81.

5. MacLeod A, Haley C, Woolliams J, Stam P. Marker densities and the mapping of ancestral junctions. Genetics Research. 2005;85(1):69–79.

6. Meuwissen T, Goddard M. Accurate prediction of genetic values for complex traits by whole-genome resequencing. Genetics. 2010;185(2):623–31.

7. Fragomeni BO, Lourenco DA, Masuda Y, Legarra A, Misztal I. Incorporation of causative quantitative trait nucleotides in single-step GBLUP. Genetics Selection Evolution. 2017;49(1):59.

8. Pérez-Enciso M, Rincón JC, Legarra A. Sequence-vs. chip-assisted genomic selection: accurate biological information is advised. Genetics Selection Evolution. 2015;47(1):1–14.

9. Fragomeni B, Lourenco D, Legarra A, VanRaden P, Misztal I. Alternative SNP weighting for single-step genomic best linear unbiased predictor evaluation of stature in US Holsteins in the presence of selected sequence variants. Journal of dairy science. 2019;102(11):10012–9.

10. Zhang C, Kemp RA, Stothard P, Wang Z, Boddicker N, Krivushin K, et al. Genomic evaluation of feed efficiency component traits in Duroc pigs using 80K, 650K and whole-genome sequence variants. Genetics Selection Evolution. 2018;50(1):1–13.

11. Veerkamp RF, Bouwman AC, Schrooten C, Calus MP. Genomic prediction using preselected DNA variants from a GWAS with whole-genome sequence data in Holstein–Friesian cattle. Genetics Selection Evolution. 2016;48(1):95.

12. Misztal I, Pocrnic I, Lourenco D, editors. Factors Influencing Accuracy of Genomic Selection with Sequence Information. JOURNAL OF ANIMAL SCIENCE; 2021: OXFORD UNIV PRESS INC JOURNALS DEPT, 2001 EVANS RD, CARY, NC 27513 USA.

13. Sargolzaei M, Schenkel FS. QMSim: a large-scale genome simulator for livestock. Bioinformatics. 2009;25(5):680–1.

14. Pocrnic I, Lourenco DA, Masuda Y, Misztal I. Accuracy of genomic BLUP when considering a genomic relationship matrix based on the number of the largest eigenvalues: a simulation study. Genetics Selection Evolution. 2019;51(1):1–10.

15. Misztal I, Tsuruta S, Lourenco D, Aguilar I, Legarra A, Vitezica Z. Manual for BLUPF90 family of programs. Athens: University of Georgia. 2014.

16. Cleveland W, Grosse E, Shyu W. Local regression models. Chapter 8 in Statistical models in S (JM Chambers and TJ Hastie eds.), 608 p. Wadsworth & Brooks/Cole, Pacific Grove, CA. 1992.

17. Zhou X, Stephens M. Genome-wide efficient mixed-model analysis for association studies. Nature genetics. 2012;44(7):821–4.

18. Wang H, Misztal I, Aguilar I, Legarra A, Muir W. Genome-wide association mapping including phenotypes from relatives without genotypes. Genetics Research. 2012;94(2):73–83.

19. Aguilar I, Misztal I, Johnson D, Legarra A, Tsuruta S, Lawlor T. Hot topic: A unified approach to utilize phenotypic, full pedigree, and genomic information for genetic evaluation of Holstein final score. Journal of Dairy Science. 2010;93(2):743–52.

20. VanRaden PM. Efficient methods to compute genomic predictions. Journal of dairy science. 2008;91(11):4414–23.

21. Zhang X, Lourenco D, Aguilar I, Legarra A, Misztal I. Weighting strategies for single-step genomic BLUP: an iterative approach for accurate calculation of GEBV and GWAS. Frontiers in genetics. 2016;7:151.

22. Lourenco D, Fragomeni B, Bradford H, Menezes I, Ferraz J, Aguilar I, et al. Implications of SNP weighting on single-step genomic predictions for different reference population sizes. Journal of Animal Breeding and Genetics. 2017;134(6):463–71.

23. Canty AJ. Resampling methods in R: the boot package. The Newsletter of the R Project Volume. 2002;2:3.

24. Moghaddar N, Khansefid M, van der Werf JH, Bolormaa S, Duijvesteijn N, Clark SA, et al. Genomic prediction based on selected variants from imputed whole-genome sequence data in Australian sheep populations. Genetics Selection Evolution. 2019;51(1):72.

25. Lopez BIM, An N, Srikanth K, Lee S, Oh J-D, Shin D-H, et al. Genomic Prediction Based on SNP Functional Annotation Using Imputed Whole-Genome Sequence Data in Korean Hanwoo Cattle. Frontiers in genetics. 2021:1523.

26. VanRaden PM, Tooker ME, O’connell JR, Cole JB, Bickhart DM. Selecting sequence variants to improve genomic predictions for dairy cattle. Genetics Selection Evolution. 2017;49(1):32.

27. MacLeod I, Bowman P, Vander Jagt C, Haile-Mariam M, Kemper K, Chamberlain A, et al. Exploiting biological priors and sequence variants enhances QTL discovery and genomic prediction of complex traits. BMC genomics. 2016;17(1):1–21.

28. de Las Heras-Saldana S, Lopez BI, Moghaddar N, Park W, Park J-e, Chung KY, et al. Use of gene expression and whole-genome sequence information to improve the accuracy of genomic prediction for carcass traits in Hanwoo cattle. Genetics Selection Evolution. 2020;52(1):1–16.

29. Pocrnic I, Lourenco DA, Masuda Y, Misztal I. Dimensionality of genomic information and performance of the Algorithm for Proven and Young for different livestock species. Genetics Selection Evolution. 2016;48(1):1–9.

30. Gozalo-Marcilla M, Buntjer J, Johnsson M, Batista L, Diez F, Werner CR, et al. Genetic architecture and major genes for backfat thickness in pig lines of diverse genetic backgrounds. Genetics Selection Evolution. 2021;53(1):1–14.

31. Baldwin-Brown JG, Long AD, Thornton KR. The power to detect quantitative trait loci using resequenced, experimentally evolved populations of diploid, sexual organisms. Molecular biology and evolution. 2014;31(4):1040–55.

32. Takeda M, Uemoto Y, Satoh M. Effect of genotyped bulls with different numbers of phenotyped progenies on quantitative trait loci detection and genomic evaluation in a simulated cattle population. Animal Science Journal. 2020;91(1):e13432.

33. van den Berg I, Fritz S, Boichard D. QTL fine mapping with Bayes C (π): a simulation study. Genetics Selection Evolution. 2013;45(1):1–11.

34. Daetwyler HD, Villanueva B, Woolliams JA. Accuracy of predicting the genetic risk of disease using a genome-wide approach. PloS one. 2008;3(10):e3395.

35. Hayes BJ, Visscher PM, Goddard ME. Increased accuracy of artificial selection by using the realized relationship matrix. Genetics research. 2009;91(1):47–60.

36. Boddhireddy P, Kelly M, Northcutt S, Prayaga K, Rumph J, DeNise S. Genomic predictions in Angus cattle: comparisons of sample size, response variables, and clustering methods for cross-validation. Journal of animal science. 2014;92(2):485–97.

37. Moser G, Tier B, Crump RE, Khatkar MS, Raadsma HW. A comparison of five methods to predict genomic breeding values of dairy bulls from genome-wide SNP markers. Genetics Selection Evolution. 2009;41(1):56.

38. Daetwyler HD, Pong-Wong R, Villanueva B, Woolliams JA. The impact of genetic architecture on genome-wide evaluation methods. Genetics. 2010;185(3):1021–31.

39. Misztal I. Inexpensive computation of the inverse of the genomic relationship matrix in populations with small effective population size. Genetics. 2016;202(2):401–9.

