## Additional file 1 for "Dimensionality of genomic information and its impact on GWA and variant selection: a simulation study"

**Table 1. Number of significantly identified QTN, SNP and the variance explained by QTN**

**(a) Ne20**

| | Ne20 Q200 H0.3 | | Ne20 Q200 H0.9 | | Ne20 Q200 H0.99 | | Ne20 Q2000 H0.3 | | Ne20 Q2000 H0.9 | | Ne20 Q2000 H0.99 | |
|---|---|---|---|---|---|---|---|---|---|---|---|---|
| | NQTN | NQTN + SNP | NQTN | NQTN + SNP | NQTN | NQTN + SNP | NQTN | NQTN + SNP | NQTN | NQTN + SNP | NQTN | NQTN + SNP |
| EIG50 | 0 (0) | 0 | 0 (0) | 0 | 0 (0) | 0 | 0 (0) | 0 | 0 (0) | 0 | 0 (0) | 0 |
| EIG60 | 0 (0) | 0 | 0 (0) | 0 | 0 (0) | 0 | 0 (0) | 0 | 0 (0) | 0 | 0 (0) | 0 |
| EIG70 | 0 (0) | 0 | 0 (0) | 0 | 1 (3.9) | 12 | 0 (0) | 0 | 0 (0) | 0 | 0 (0) | 0 |
| EIG80 | 0 (0) | 0 | 1 (10.4) | 23 | 1 (8) | 16 | 0 (0) | 0 | 0 (0) | 0 | 0 (0) | 0 |
| EIG90 | 1 (9.8) | 10 | 3 (17.6) | 95 | 6 (21.4) | 133 | 0 (0) | 0 | 0 (0) | 0 | 0 (0) | 0 |
| EIG95 | 2 (15.3) | 20 | 11 (41.5) | 269 | 22 (46) | 518 | 0 (0) | 0 | 0 (0) | 0 | 3 (1.5) | 17 |
| EIG98 | 4 (24.5) | 60 | 19 (55.8) | 487 | 35 (59.9) | 1063 | 0 (0) | 0 | 7 (5.7) | 72 | 17 (11.2) | 108 |
| EIG99 | 5 (30.8) | 125 | 23 (61.5) | 639 | 49 (68) | 1345 | 0 (0) | 1 | 13 (9.9) | 99 | 33 (18.9) | 204 |
| All | 10 (44.9) | 563 | 40 (72.9) | 1208 | 75 (74.3) | 1622 | 3 (3.9) | 16 | 54 (26.8) | 378 | 142 (43.1) | 844 |

*Values in the bracket are the %Variance explained by identified QTN

**(b) Ne200**

| | Ne200<br>Q200 H0.3 | | Ne200<br>Q200 H0.9 | | Ne200<br>Q200 H0.99 | | Ne200<br>Q2000 H0.3 | | Ne200<br>Q2000 H0.9 | | Ne200<br>Q2000 H0.99 | |
|---|---|---|---|---|---|---|---|---|---|---|---|---|
| | NQTN | NQTN + SNP | NQTN | NQTN + SNP | NQTN | NQTN + SNP | NQTN | NQTN + SNP | NQTN | NQTN + SNP | NQTN | NQTN + SNP |
| EIG50 | 0 (0) | 1 | 2 (6.4) | 3 | 2 (16.5) | 14 | 0 (0) | 0 | 0 (0) | 1 | 0 (0) | 4 |
| EIG60 | 1 (8.6) | 1 | 6 (18.1) | 14 | 5 (28.2) | 39 | 0 (0) | 0 | 1 (2.3) | 1 | 0 (0) | 1 |
| EIG70 | 3 (22.3) | 5 | 14 (40.1) | 80 | 15 (48.9) | 137 | 0 (0) | 0 | 3 (4.5) | 16 | 2 (3.4) | 6 |
| EIG80 | 7 (38.2) | 29 | 29 (61.6) | 244 | 28 (65.9) | 346 | 1 (0.4) | 4 | 4 (4.7) | 15 | 8 (10.3) | 41 |
| EIG90 | 10 (48.7) | 117 | 46 (76.1) | 522 | 55 (87) | 704 | 0 (0) | 0 | 19 (17.6) | 80 | 28 (20.7) | 161 |
| EIG95 | 15 (59.3) | 201 | 57 (83) | 731 | 67 (91.8) | 989 | 0 (0) | 1 | 46 (33.2) | 199 | 71 (37.9) | 386 |
| EIG98 | 22 (68) | 297 | 75 (89.1) | 983 | 80 (95.2) | 1201 | 7 (6.8) | 11 | 74 (42.3) | 432 | 134 (52.5) | 741 |
| EIG99 | 28 (73.1) | 417 | 81 (91.3) | 1114 | 83 (95.6) | 1324 | 12 (11.8) | 30 | 92 (46.6) | 598 | 179 (59.7) | 942 |
| All | 33 (77.4) | 512 | 87 (93) | 1213 | 86 (96) | 1348 | 15 (13.9) | 53 | 109 (49.6) | 705 | 223 (65.1) | 1092 |

*Values in the bracket are the %Variance explained by identified QTN

**Table 2. Approximated sample size using local polynomial regression ($\text{Sample}_{\text{app1}}$) and proposed equation ($\text{Sample}_{\text{app2}}$) for all feasible scenarios**

**(a) Ne20 Q200**

| Heritability | %Var[1] | $SS_{pol}$[2] | $EIGx_{app1}$[3] | $SS_{rel}$[4] | $EIGx_{app2}$[5] | Diff[6] |
|---|---|---|---|---|---|---|
| 0.3 | 44.9 | 30000 (ALL) | | | | |
| 0.9 | 44.9 | 2223 | EIG95~98 (1770~4100) | 2698 | EIG95~98 (1770~4100) | 475 |
| 0.99 | 44.9 | 1662 | EIG90~95 (890~1800) | 2453 | EIG95~98 (1800~4100) | 790 |
| 0.3 | 30.8 | 6900 (EIG99) | | | | |
| 0.9 | 30.8 | 1324 | EIG90~95 (900~1770) | 646 | EIG80~90 (410~900) | -677 |
| 0.99 | 30.8 | 1181 | EIG90~95 (890~1800) | 588 | EIG80~90 (410~890) | -593 |
| 0.3 | 24.5 | 4000 (EIG98) | | | | |
| 0.9 | 24.5 | 1102 | EIG90~95 (900~1770) | 382 | EIG70~80 (240~410) | -720 |
| 0.99 | 24.5 | 966 | EIG90~95 (890~1800) | 347 | EIG70~80 (230~410) | -618 |
| 0.3 | 15.3 | 1700 (EIG95) | | | | |
| 0.9 | 15.3 | 742 | EIG80~90 (410~900) | 167 | EIG60~70 (140~240) | -575 |
| 0.99 | 15.3 | 645 | EIG80~90 (410~890) | 152 | EIG60~70 (140~230) | -493 |
| 0.3 | 9.8 | 860 (EIG90) | | | | |
| 0.9 | 9.8 | 463 | EIG70~80 (240~410) | 85 | EIG50~60 (80~140) | -377 |
| 0.99 | 9.8 | 450 | EIG80~90 (410~890) | 77 | ~EIG50 (~80) | -373 |

**(b) Ne20 Q2000**

| Heritability | %Var[1] | $SS_{pol}$[2] | $EIGx_{app1}$[3] | $SS_{rel}$[4] | $EIGx_{app2}$[5] | Diff[6] |
|---|---|---|---|---|---|---|
| 0.3 | 3.9 | 30000 (ALL) | | | | |

| | | | | | | |
|---|---|---|---|---|---|---|
| 0.9 | 3.9 | 3049 | EIG95~98 (1750~4100) | 3007 | EIG95~98 (1750~4100) | -41 |
| 0.99 | 3.9 | 2378 | EIG95~98 (1700~4000) | 2734 | EIG95~98 (1700~4000) | -356 |

**(c) Ne200 Q200**

| Heritability | %Var[1] | $SS_{pol}$[2] | $EIGx_{app1}$[3] | $SS_{rel}$[4] | $EIGx_{app2}$[5] | Diff[6] |
|---|---|---|---|---|---|---|
| 0.3 | 77.4 | 30000 (ALL) | | | | |
| 0.9 | 77.4 | 5205 | EIG80~90 (2650~5250) | 4324 | EIG80~90 (2650~5250) | -880 |
| 0.99 | 77.4 | 3814 | EIG80~90 (2700~5300) | 3931 | EIG80~90 (2700~5300) | 117 |
| 0.3 | 73.1 | 21000 (EIG99) | | | | |
| 0.9 | 73.1 | 4401 | EIG80~90 (2650~5250) | 3047 | EIG80~90 (2650~5250) | -1353 |
| 0.99 | 73.1 | 3259 | EIG80~90 (2700~5300) | 2770 | EIG80~90 (2700~5300) | -489 |
| 0.3 | 68 | 15000 (EIG98) | | | | |
| 0.9 | 68 | 3530 | EIG80~90 (2650~5250) | 2194 | EIG70~80 (1550~2650) | -1336 |
| 0.99 | 68 | 2600 | EIG70~80 (1600~2700) | 1995 | EIG70~80 (1600~2700) | -605 |
| 0.3 | 59.3 | 8600 (EIG95) | | | | |
| 0.9 | 59.3 | 2455 | EIG70~80 (1550~2650) | 1277 | EIG60~70 (910~1550) | -1178 |
| 0.99 | 59.3 | 2178 | EIG70~80 (1600~2700) | 1161 | EIG60~70 (940~1600) | -1017 |
| 0.3 | 48.7 | 5100 (EIG90) | | | | |
| 0.9 | 48.7 | 1980 | EIG70~80 (1550~2650) | 772 | EIG50~60 (520~910) | -1208 |
| 0.99 | 48.7 | 1664 | EIG70~80 (1600~2700) | 701 | EIG50~60 (550~940) | -963 |
| 0.3 | 38.2 | 2600 (EIG80) | | | | |
| 0.9 | 38.2 | 1511 | EIG60~70 (910~1550) | 401 | ~EIG50 (~520) | -1110 |
| 0.99 | 38.2 | 1235 | EIG60~70 (940~1600) | 365 | ~EIG50 (~550) | -870 |

| | | | | | | |
|---|---|---|---|---|---|---|
| 0.3 | 22.3 | 1500 (EIG70) | | | | |
| 0.9 | 22.3 | 984 | EIG60~70 (910~1550) | 237 | ~EIG50 (~520) | -747 |
| 0.99 | 22.3 | 676 | EIG50~60 (550~940) | 215 | ~EIG50 (~550) | -461 |
| 0.3 | 8.6 | 900 (EIG60) | | | | |
| 0.9 | 8.6 | 583 | EIG50~60 (520~910) | 138 | ~EIG50 (~520) | -445 |
| 0.99 | 8.6 | NA | NA | 125 | ~EIG50 (~550) | NA |

**(d) Ne200 Q2000**

| Heritability | %Var[1] | $SS_{pol}$[2] | $EIGx_{app1}$[3] | $SS_{rel}$[4] | $EIGx_{app2}$[5] | Diff[6] |
|---|---|---|---|---|---|---|
| 0.3 | 13.9 | 30000 (ALL) | | | | |
| 0.9 | 13.9 | 4524 | EIG80~EIG90 (2600~5100) | 4719 | EIG80~EIG90 (2600~5100) | 194 |
| 0.99 | 13.9 | 3664 | EIG80~EIG90 (2600~5100) | 4290 | EIG80~EIG90 (2600~5100) | 625 |
| 0.3 | 11.8 | 22000 (EIG99) | | | | |
| 0.9 | 11.8 | 4032 | EIG80~EIG90 (2600~5100) | 3437 | EIG80~EIG90 (2600~5100) | -595 |
| 0.99 | 11.8 | 3176 | EIG80~EIG90 (2600~5100) | 3124 | EIG80~EIG90 (2600~5100) | -52 |
| 0.3 | 6.8 | 15200 (EIG98) | | | | |
| 0.9 | 6.8 | 2860 | EIG80~EIG90 (2600~5100) | 2292 | EIG70~EIG80 (1500~2600) | -569 |
| 0.99 | 6.8 | 2088 | EIG70~EIG80 (1500~2600) | 2084 | EIG70~EIG80 (1500~2600) | -4 |

*%Var1: percentage of variance explained by significantly identified QTN

*$SS_{pol}$2: Approximated sample size using local polynomial regression

*$EIGx_{app1}$3: EIGx scenario range including $Sample_{app1}$

*$SS_{rel}$4: Approximated sample size using proposed equation

*$EIGx_{app2}$5: EIGx scenario range including $Sample_{app2}$

*Diff6: Difference between $Sample_{app2}$ and $Sample_{app1}$

**Figure 1. GWAS results for all scenarios**

**(a) Ne20 Q200 H0.3**

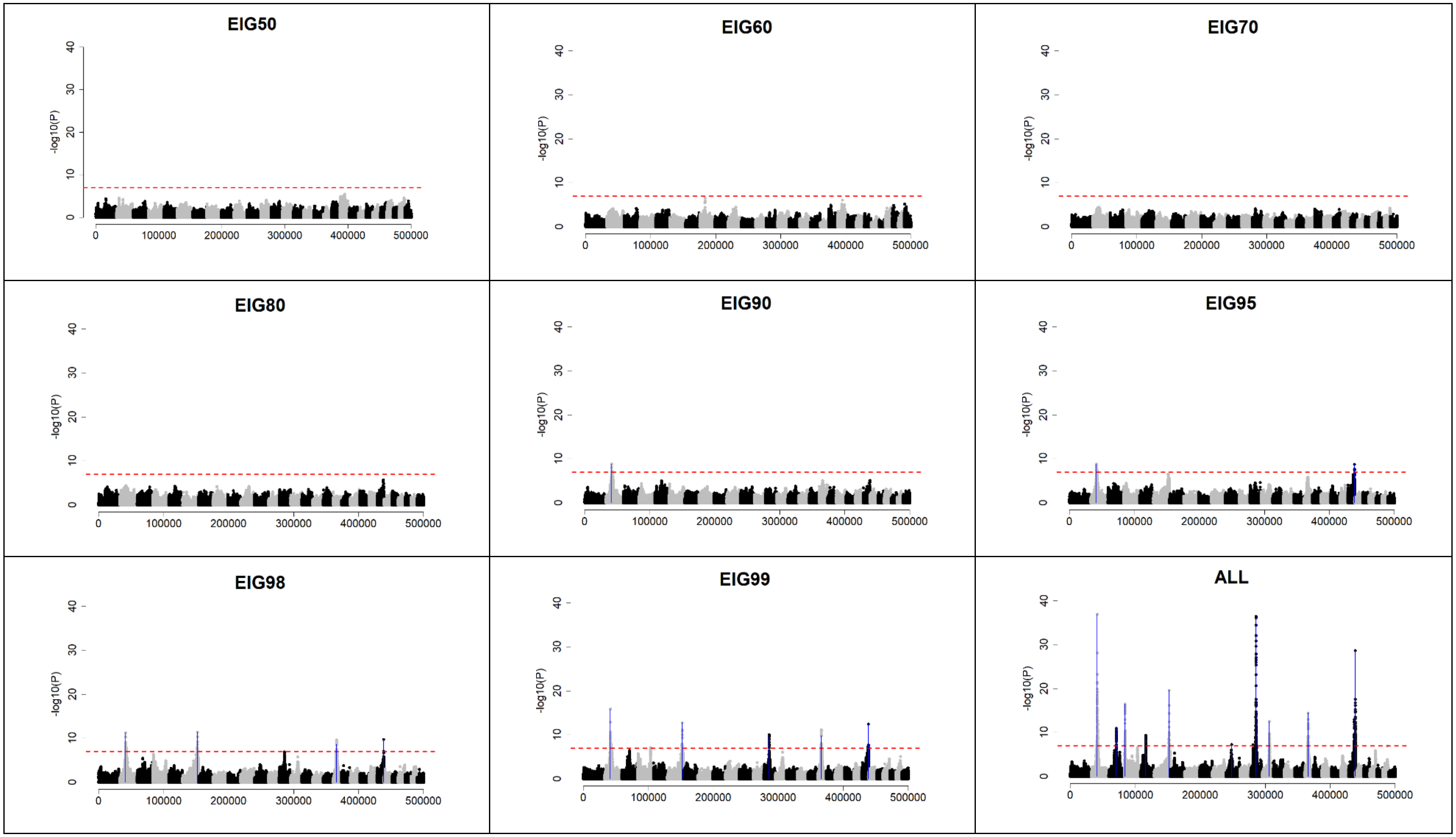

**(b) Ne20 Q200 H0.9**

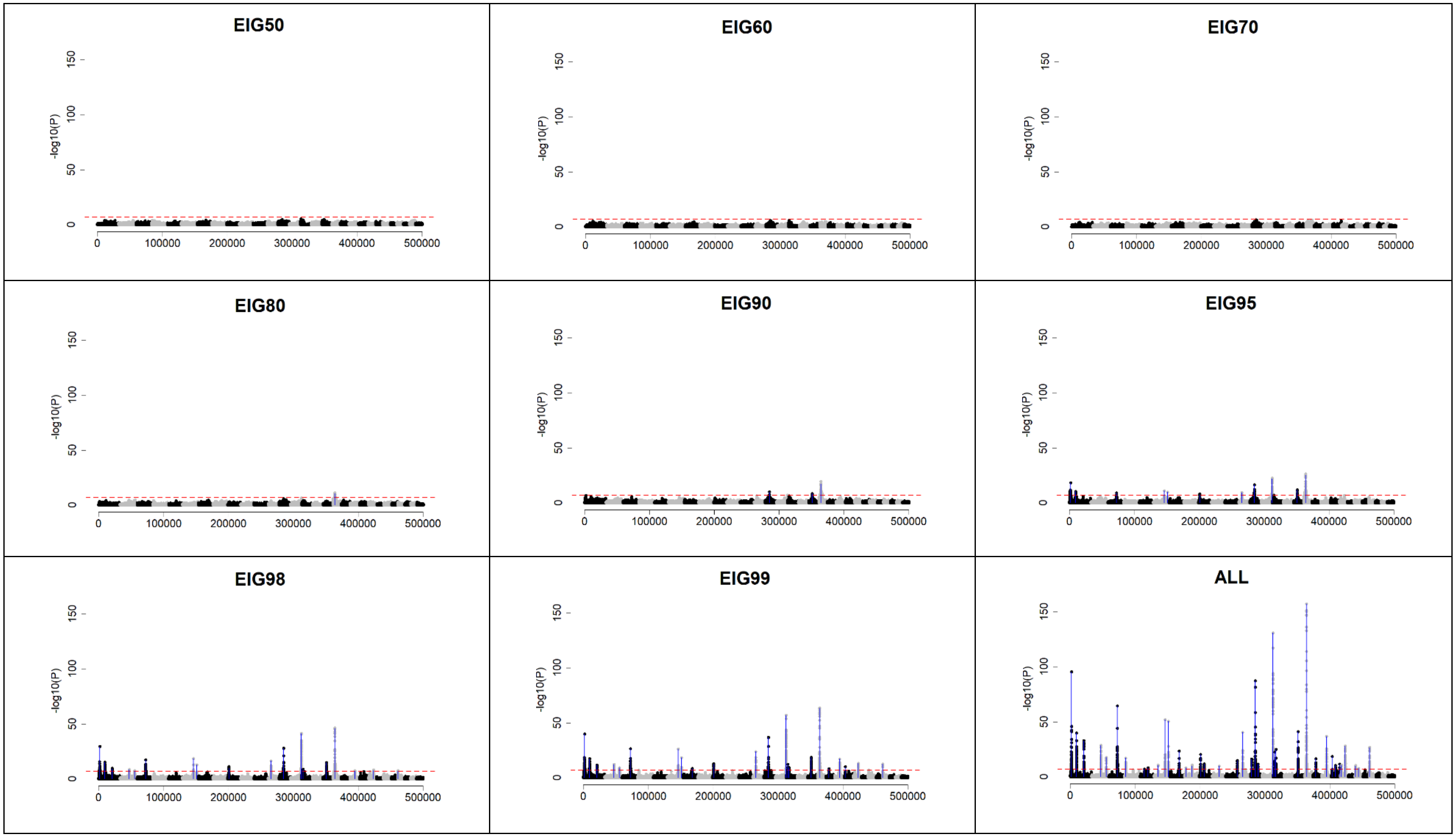

## (c) Ne20 Q200 H0.99

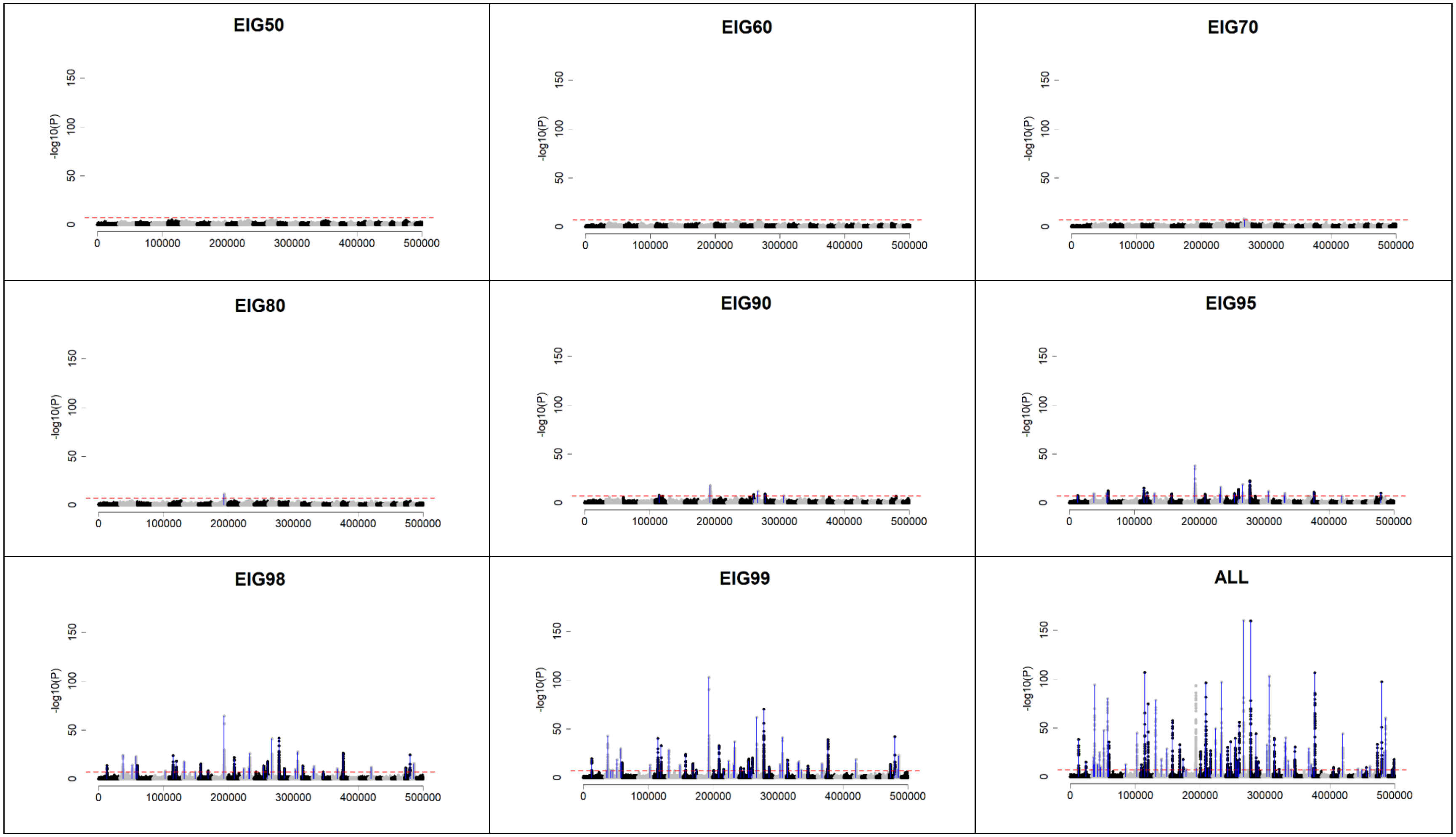

**(d) Ne20 Q2000 H0.3**

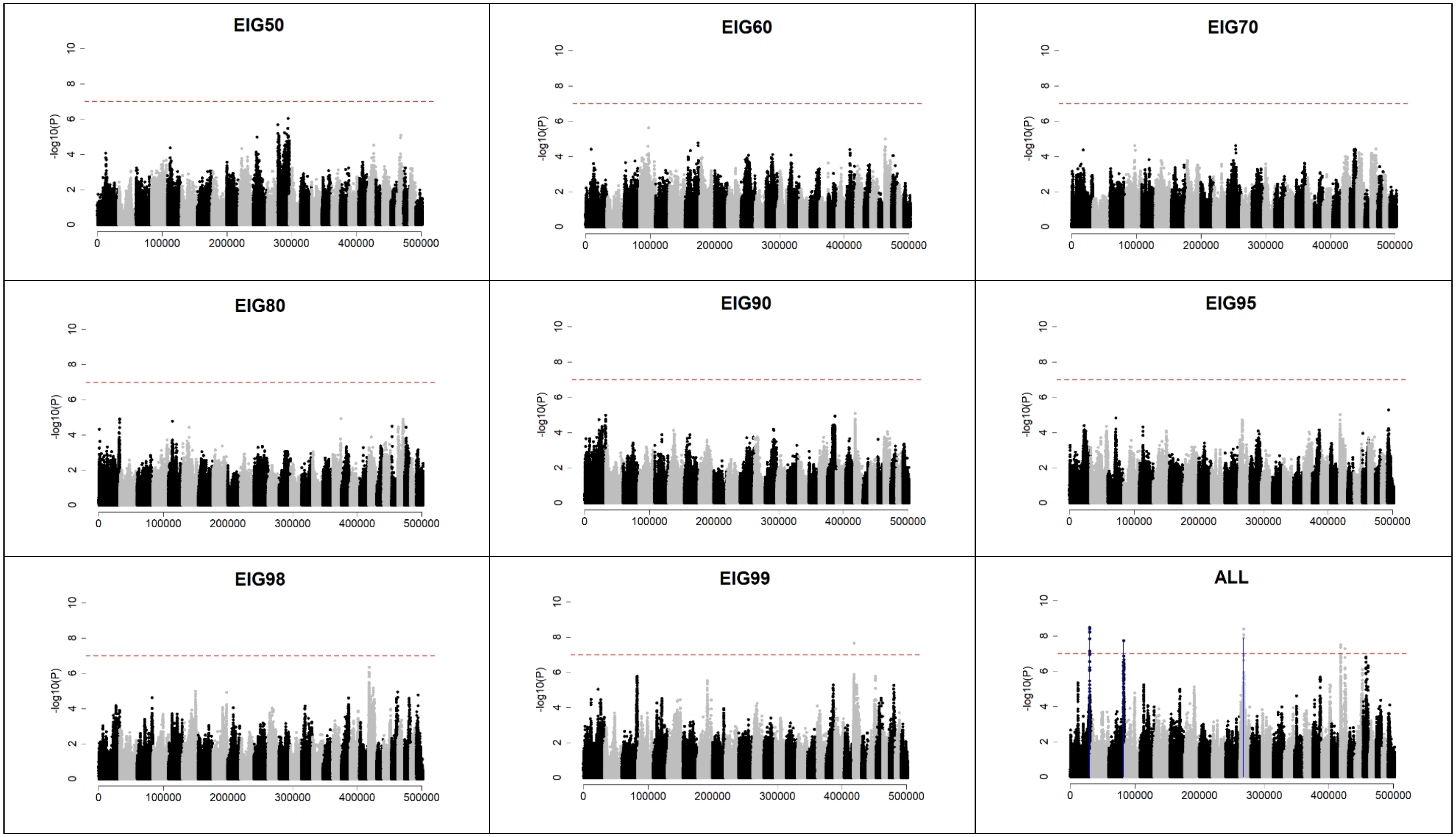

(e) Ne20 Q2000 H0.9

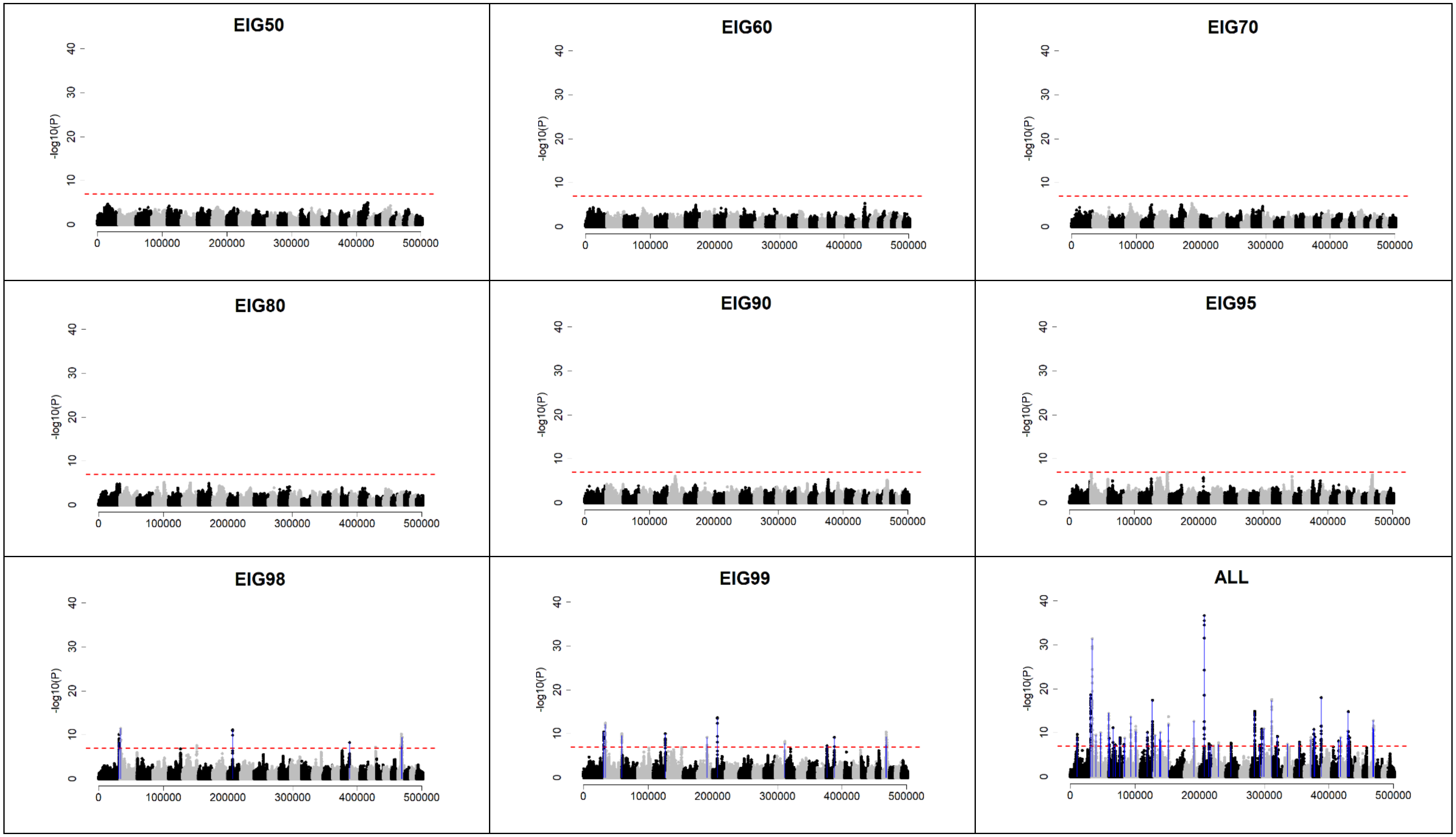

**(f) Ne20 Q2000 H0.99**

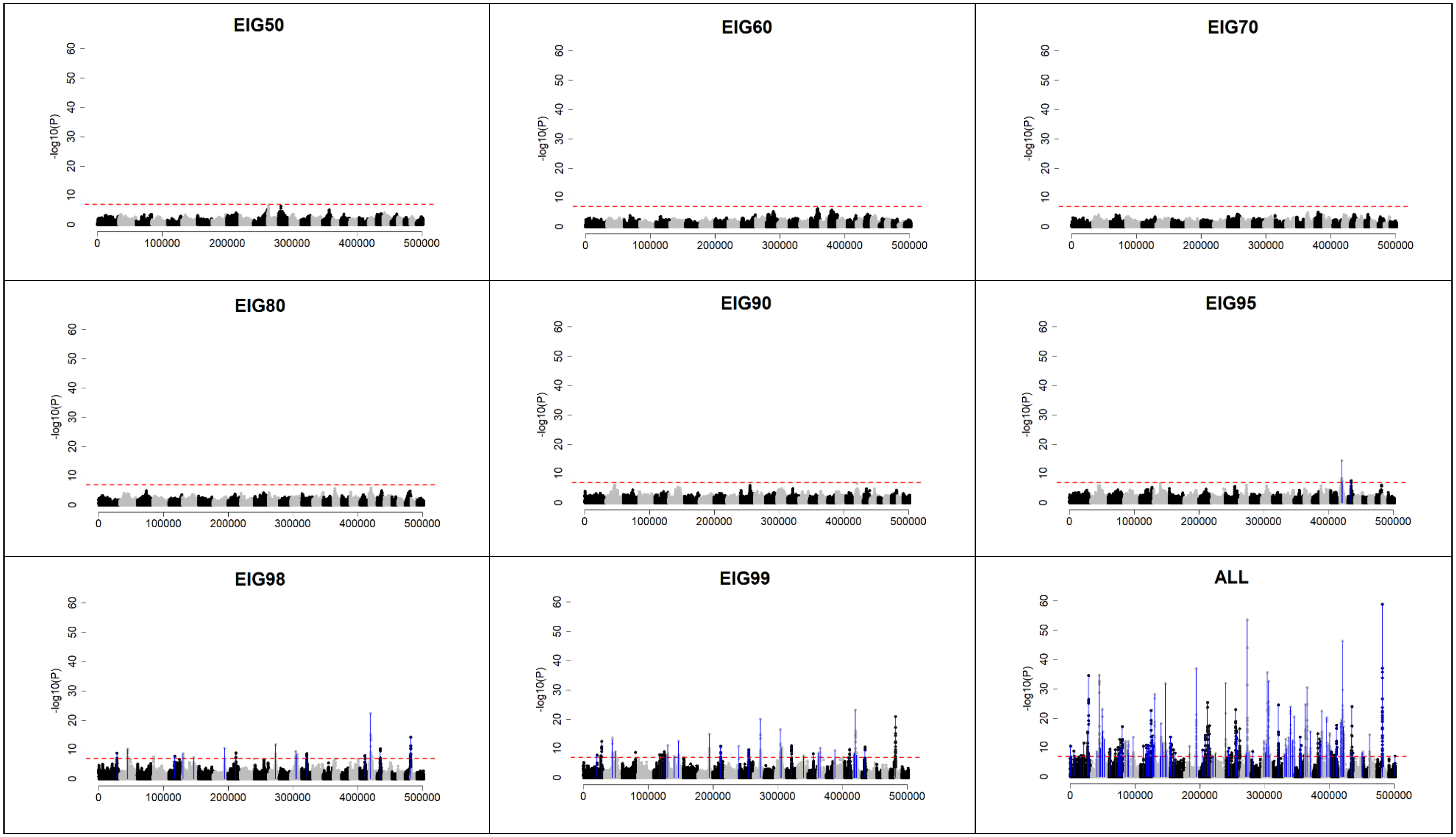

(g) Ne200 Q200 H0.3

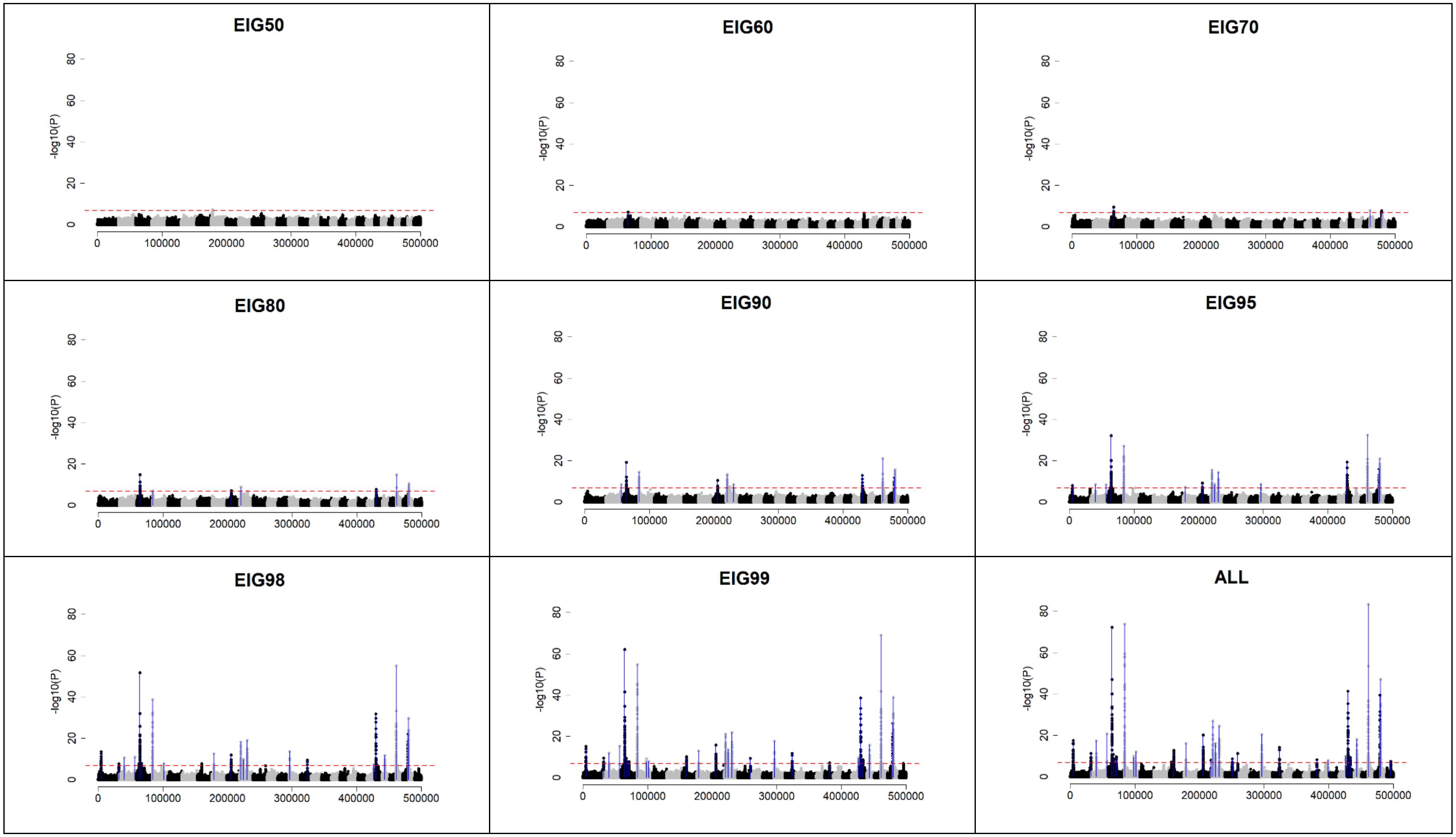

(h) Ne200 Q200 H0.9

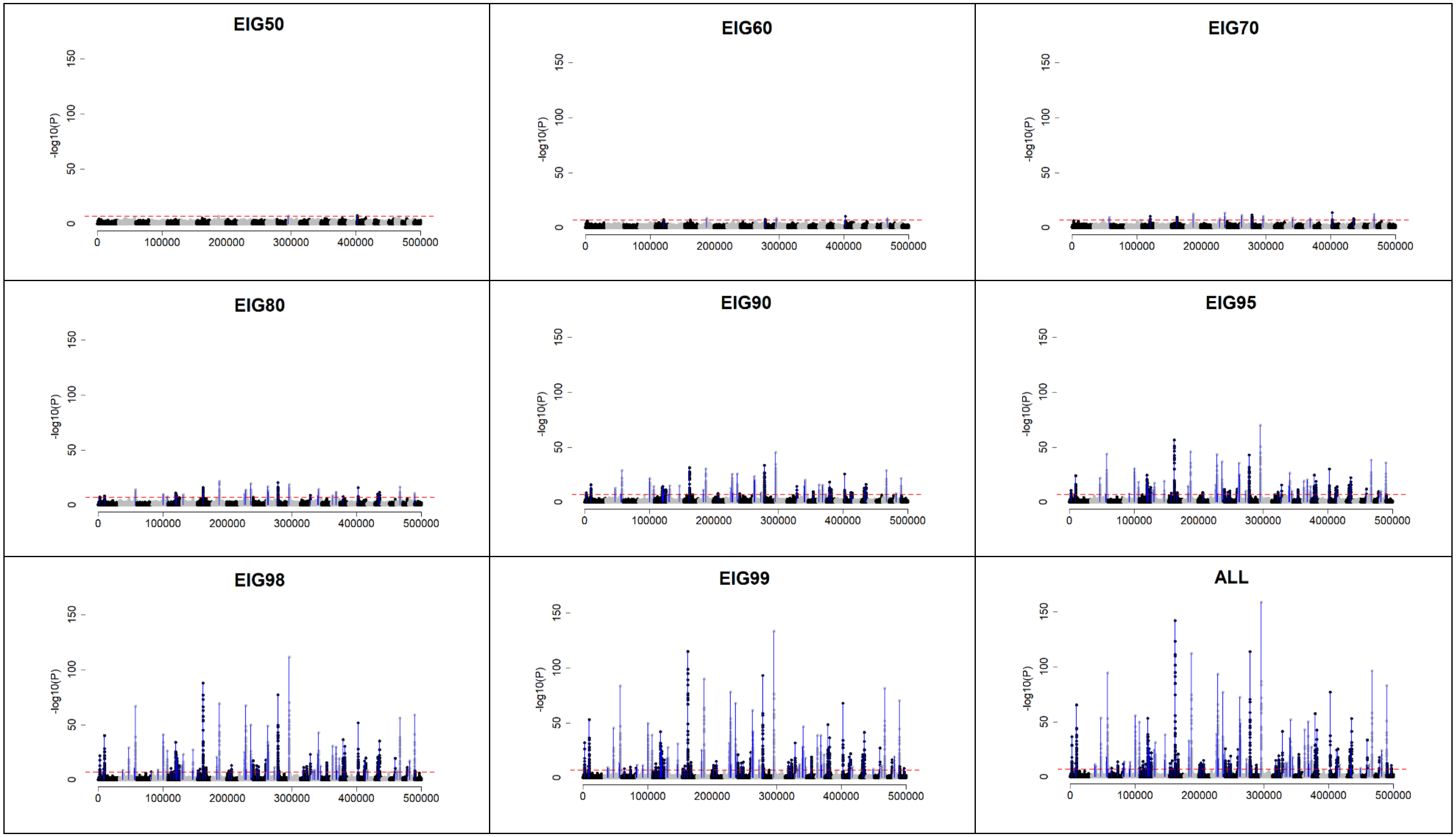

## (i) Ne200 Q200 H0.99

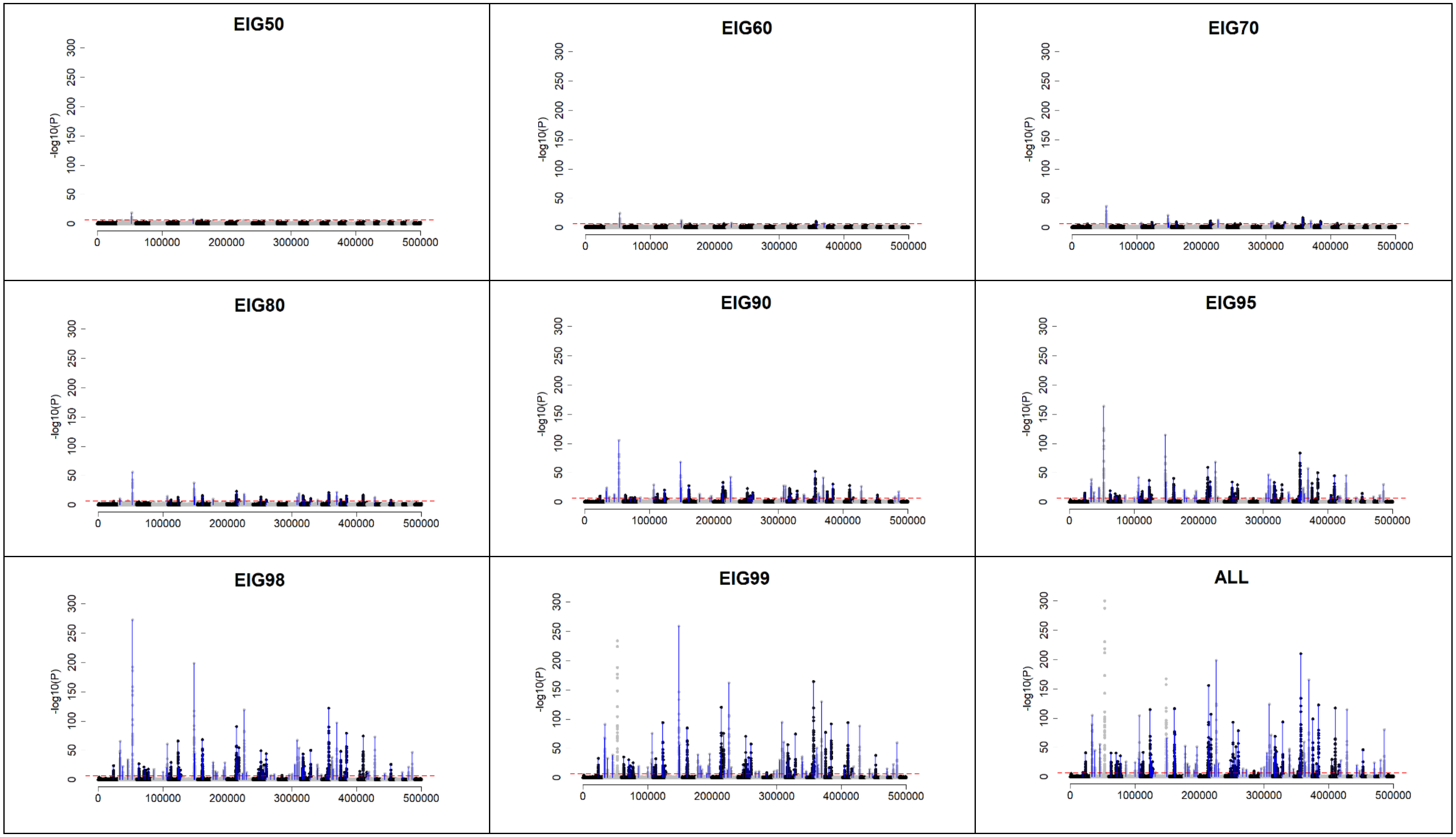

(j) Ne200 Q2000 H0.3

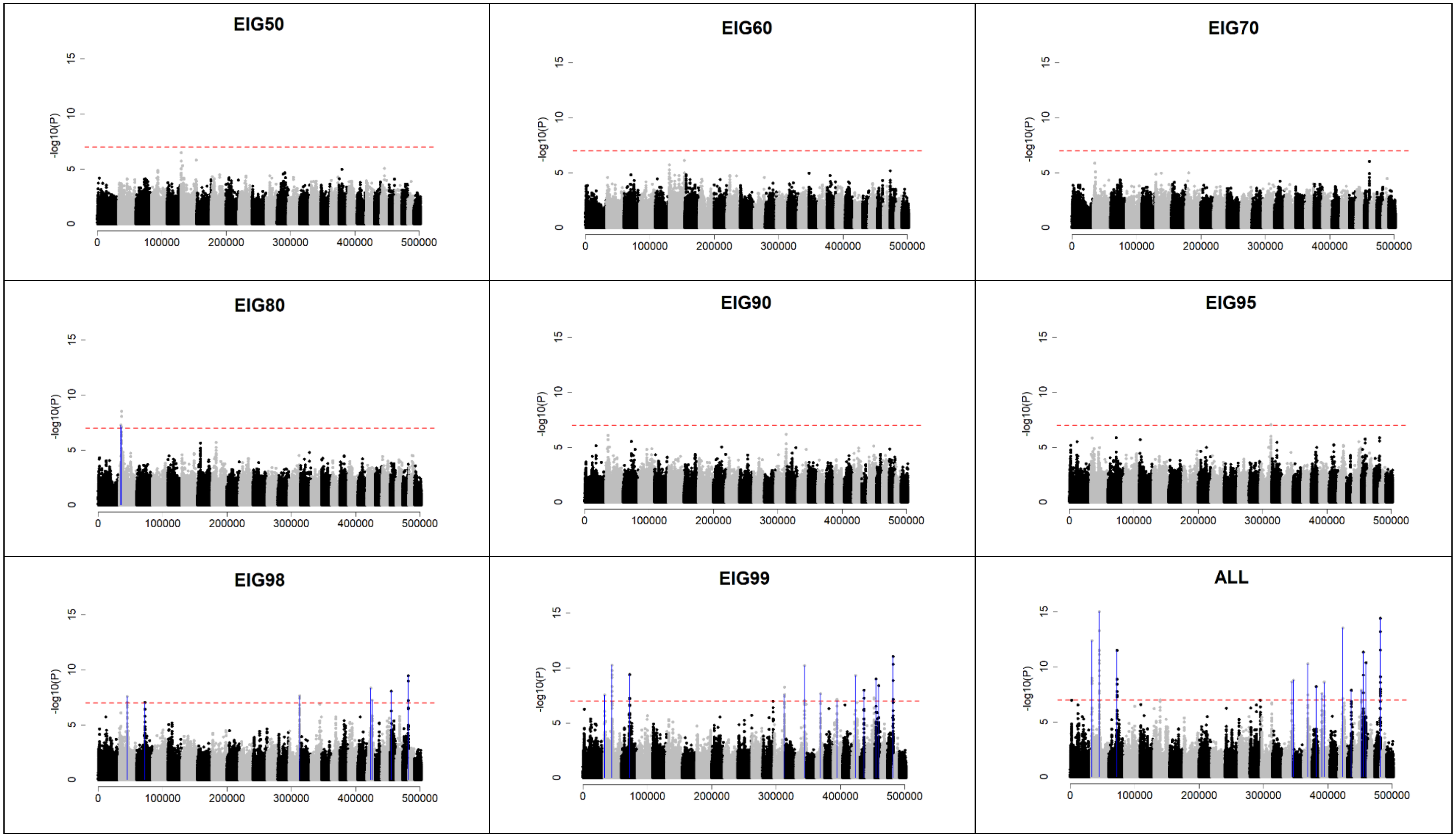

(k) Ne200 Q2000 H0.9

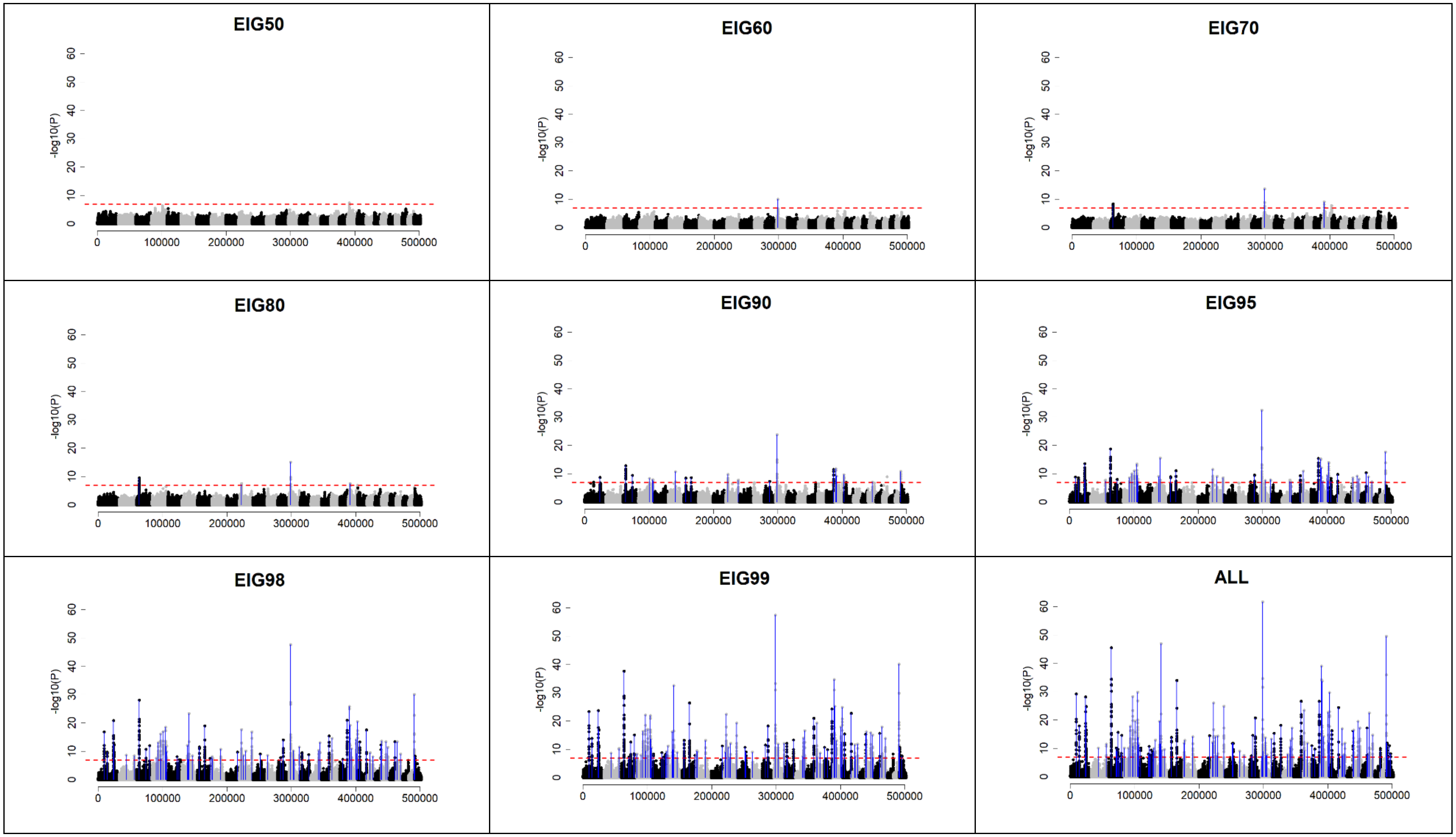

## (l) Ne200 Q2000 H0.99

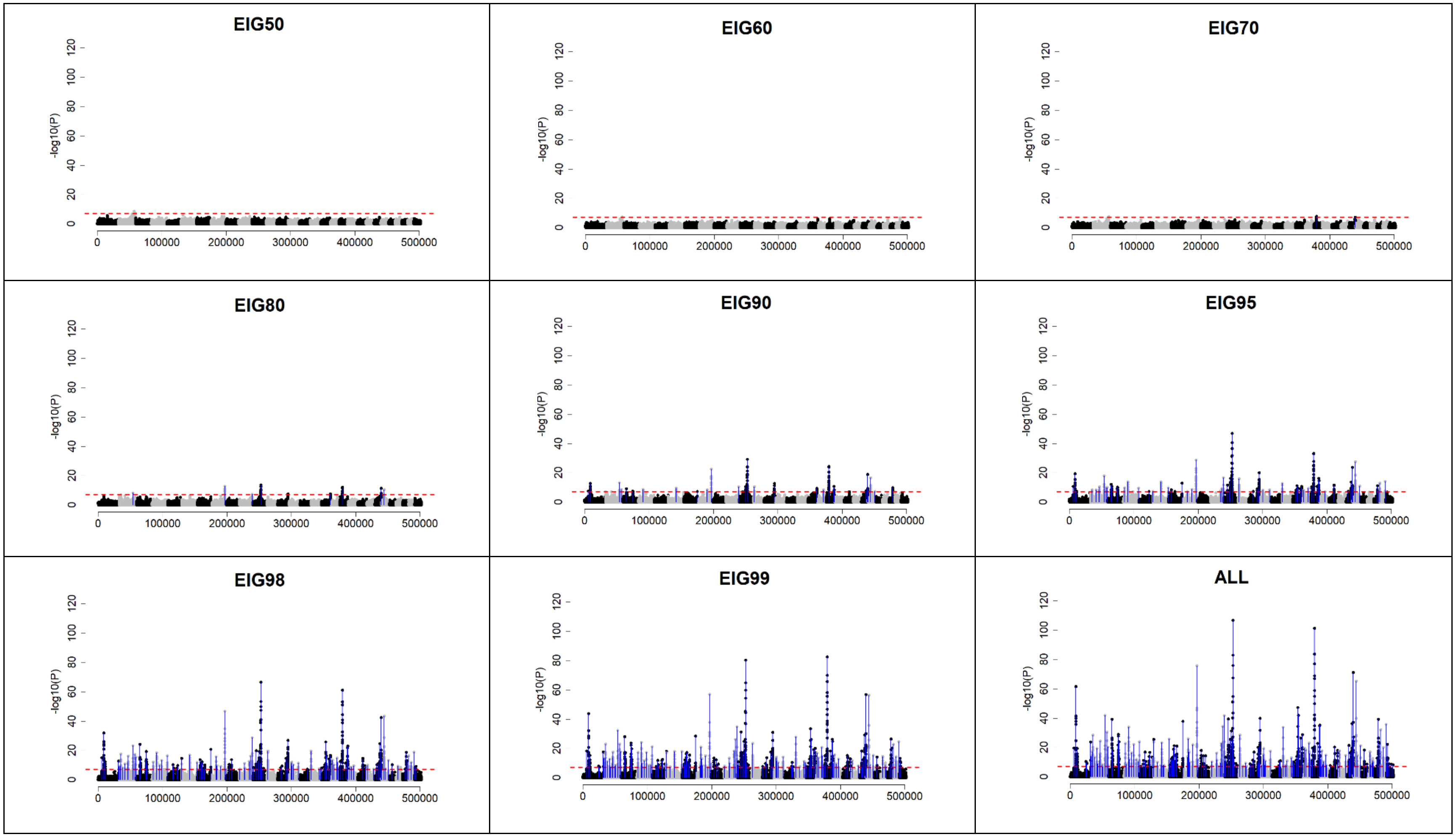

**Figure 2. Prediction accuracy when only QTN (N = 200, 2000 for Q200, Q2000 scenarios) and those QTN were added to 50k under H99 scenario**

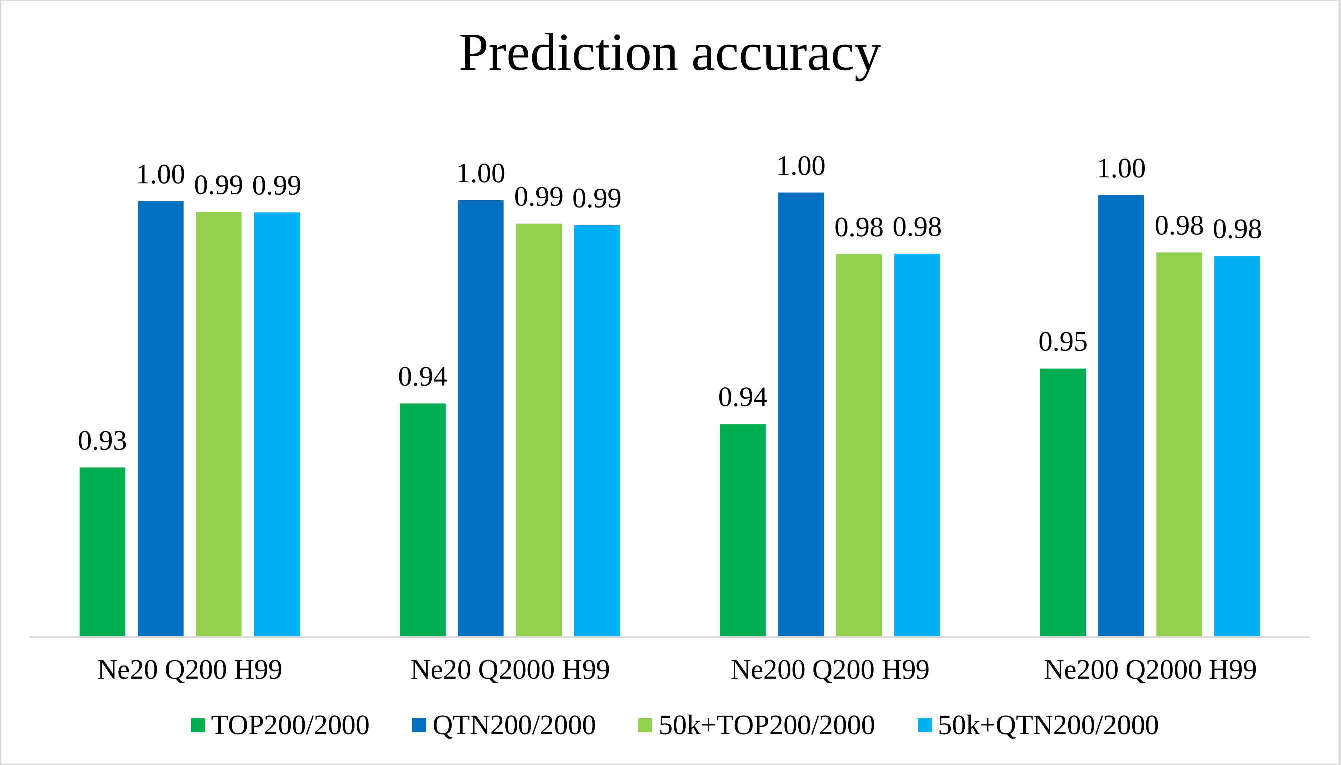
